# Multiple sclerosis iPSC-derived oligodendroglia conserve their intrinsic properties to functionally interact with axons and glia in vivo

**DOI:** 10.1101/2020.05.05.078642

**Authors:** Sabah Mozafari, Laura Starost, Blandine Manot-Saillet, Beatriz Garcia-Diaz, Yu Kang T. Xu, Delphine Roussel, Marion J. F. Levy, Linda Ottoboni, Kee-Pyo Kim, Hans R. Schöler, Timothy E. Kennedy, Jack P. Antel, Gianvito Martino, Maria Cecilia Angulo, Tanja Kuhlmann, Anne Baron-Van Evercooren

## Abstract

The remyelination failure in multiple sclerosis (MS) is associated with a migration/differentiation block of oligodendroglia. The reason for this block is highly debated. It could result from disease-related extrinsic regulators of the oligodendroglial biology or reflect MS oligodendrocyte intrinsic properties. To avoid confounding immune-mediated extrinsic effect, we used an immune-deficient, dysmyelinating mouse model, to compare side-by-side induced pluripotent stem-cell-derived O4+ oligodendroglia from MS and healthy donors following their engraftment in the developing CNS. We show that the MS-progeny survives, proliferates and differentiates into oligodendrocytes to the same extent as controls. Quantitative multi-parametric imaging indicates that MS and control oligodendrocytes generate equal amounts of myelin, with *bona-fide* nodes of Ranvier and promote equal restoration of their host slow conduction. Moreover, the MS-derived progeny expressed oligodendrocyte- and astrocyte-specific connexins and established functional connections with donor and host glial cells. Thus, MS pluripotent stem cell-derived progeny fully integrates into functional axo-glial and glial-glial components, reinforcing the view that the MS oligodendrocyte differentiation block is not due to intrinsic oligodendroglial deficits. These biological findings as well as the fully integrated human-murine chimeric model should facilitate the development of pharmacological or cell-based therapies to promote CNS remyelination.

**One Sentence Summary:** Multiple Sclerosis oligodendroglia, regardless of major immune manipulators, are intrinsically capable of myelination and making functional axo-glia and glia-glia connections after engraftment in the murine CNS, reinforcing the view that the MS oligodendrocyte differentiation block is not due to major intrinsic oligodendroglial deficits but most likely to environmental conditions.

## Introduction

Remyelination occurs in multiple sclerosis (MS) lesions but its capacity decreases over time (*1-5*). Failed remyelination in MS leads to altered conduction followed by axon degeneration, which in the long run, results in severe and permanent neurological deficits (*6*). MS lesions may or may not harbor immature oligodendroglia (oligodendrocyte progenitors and pre-oligodendrocytes), with such cells failing to differentiate into myelin-forming cells, suggesting that oligodendrocyte differentiation is blocked (*7-9*). So far, the mechanism underlying this block is poorly understood. It may result from adverse environmental conditions or the failed capacity of oligodendrocyte progenitors/pre-oligodendrocytes to migrate or mature efficiently into myelin-forming cells or even a combination of these conditions. It has been shown that increasing remyelination either through manipulating the endogenous pool (*10, 11*) or by grafting competent myelin forming oligodendroglia (*12, 13*) or both (*14*) can restore the lost axonal functions, improve the clinical score and might protect from subsequent axonal degeneration in experimental models (*15, 16*) or clinical studies (*5*).

There are multiple ways to investigate the oligodendroglial lineage in disease. Cells can be studied in post-mortem tissue sections or purified from post-mortem adult human brain for in vitro and transcriptomic/proteomic analysis. In this respect, in vitro experiments highlighted the heterogeneity of the adult human oligodendrocyte progenitor population in terms of antigen and miRNA expression, suggesting that remyelination in the adult human brain involves multiple populations of progenitors (*17*). Moreover, single cell transcriptomics characterized in detail the heterogeneity of human oligodendroglial cells, emphasizing changes in MS, with some sub-populations expressing disease specific markers that could play a role in disease onset and/or aggravation (*18, 19*).

Yet this MS signature could pre-exist or be acquired early at disease onset. Moreover, most of these MS post-mortem analyses or experimental models cannot overlook the involvement of extrinsic factors such as immune factors that might add more complexity towards understanding the behavior of MS oligodenroglial cells.

Little is known about the biology of the MS oligodendroglial lineage, primarily due to the impossibility, for ethical reasons, to harvest oligodendroglial populations from patients and study the diseased cells and their matching controls in vitro or in vivo, after cell transplantation. While cell-cell interactions and cell heterogeneity in diseased conditions generate more complexity when comparing control and pathological samples, the induced pluripotent stem cell (iPSC) technology provides a unique opportunity to study homogeneous populations of human oligodendroglial cells and gain further insights into monogenetic diseases and multi-factorial diseases, such as MS. Indeed, the iPSC technology has unraveled differences in oligodendroglia biology, in Huntington’s disease (*20*) and schizophrenia (*21, 22*) indicating that these cells can contribute autonomously to multifactorial diseases outcome. However, so far, little is known about the potential contribution of MS oligodendroglia to failed remyelination. While senescence affects iPSC-NPCs derived from primary progressive MS (PPMS) patients (*23, 24*), only few preliminary reports alluded to the fate of PPMS (*25, 26*) or RRMS (*27*) iPSC-derived oligodendroglia after experimental transplantation but did not study *per se* their capacity to differentiate into functional myelin-forming cells. We exploited a robust approach (*28*) to generate large quantities of O4+ oligodendroglial cells (hiOLs) from skin fibroblasts of 3 relapsing-remitting MS (RRMS) and 3 healthy subjects, including two monozygous twin pairs discordant for the disease. As a critical feature of the pluripotent-derived cells should be their ability to fully integrate and function in vivo, we compared the capacity of healthy and MS-hiOL derivatives to integrate and restore axo-glial and glial-glial functional interaction in vivo after engraftment in the developing dysmyelinated murine CNS. Our data show that in non-inflammatory conditions, the intrinsic properties of oligodendroglial cells to differentiate, myelinate and establish functional cell-cell interactions in vivo are not altered in MS and can make them candidates of interest for personalized drug/cell therapies as pluripotency maintains MS oliogdendroglial cells in a genuine “non pathological” state.

## Results

Fibroblasts were isolated from 3 controls and 3 MS patients and reprogrammed into iPSC. Pluripotent cells were differentiated into NPCs and further into O4+ hiOLs for 12 days in vitro under GDM conditions as previously described (*28*). All hiOL lines were fully characterized in vitro. hiOLs from all lines were selected using flow cytometry for O4 before transplantation. Since our aim was to study the intrinsic properties of MS cells, we chose to engraft O4+ hiOLs in the purely dysmyelinating Shi/Shi:Rag2-/- mouse model to avoid confounding immune-mediated extrinsic effects.

### MS hiOL do not exhibit aberrant survival or proliferation in vivo

We first questioned whether MS-hiOLs differed from control-hiOLs (WT) in their capacity to survive and proliferate in vivo. To this aim, we grafted MS- and control-hiOLs in the forebrain of neonatal Shi/Shi:Rag2-/- mice. MS cells engrafted (1 injection/hemisphere) in the rostral forebrain, spread primarily through white matter, including the corpus callosum and fimbria, as previously observed using human fetal progenitors (*13, 29, 30*) and control-hiOLs (*28, 31*). With time, cells also spread rostrally to the olfactory bulb, and caudally to the brain stem and cerebellum (fig. S1). Examining engrafted brains at 8, 12 and 16 weeks post graft (wpg), we found that MS-hiOLs expressing the human nuclear marker STEM101 and the oligodendroglial specific transcription factor OLIG2, maintained a slow proliferation rate at all times (5-19% of STEM+ cells), with no difference in Ki67+ MS-hiOLs compared to control (Fig. 1A and C). Moreover, immunostaining for cleaved Caspase-3 at 8 wpg indicated that MS cells survived equally as well as control-hiOLs (Fig. 1B and D).

**Fig. 1.**
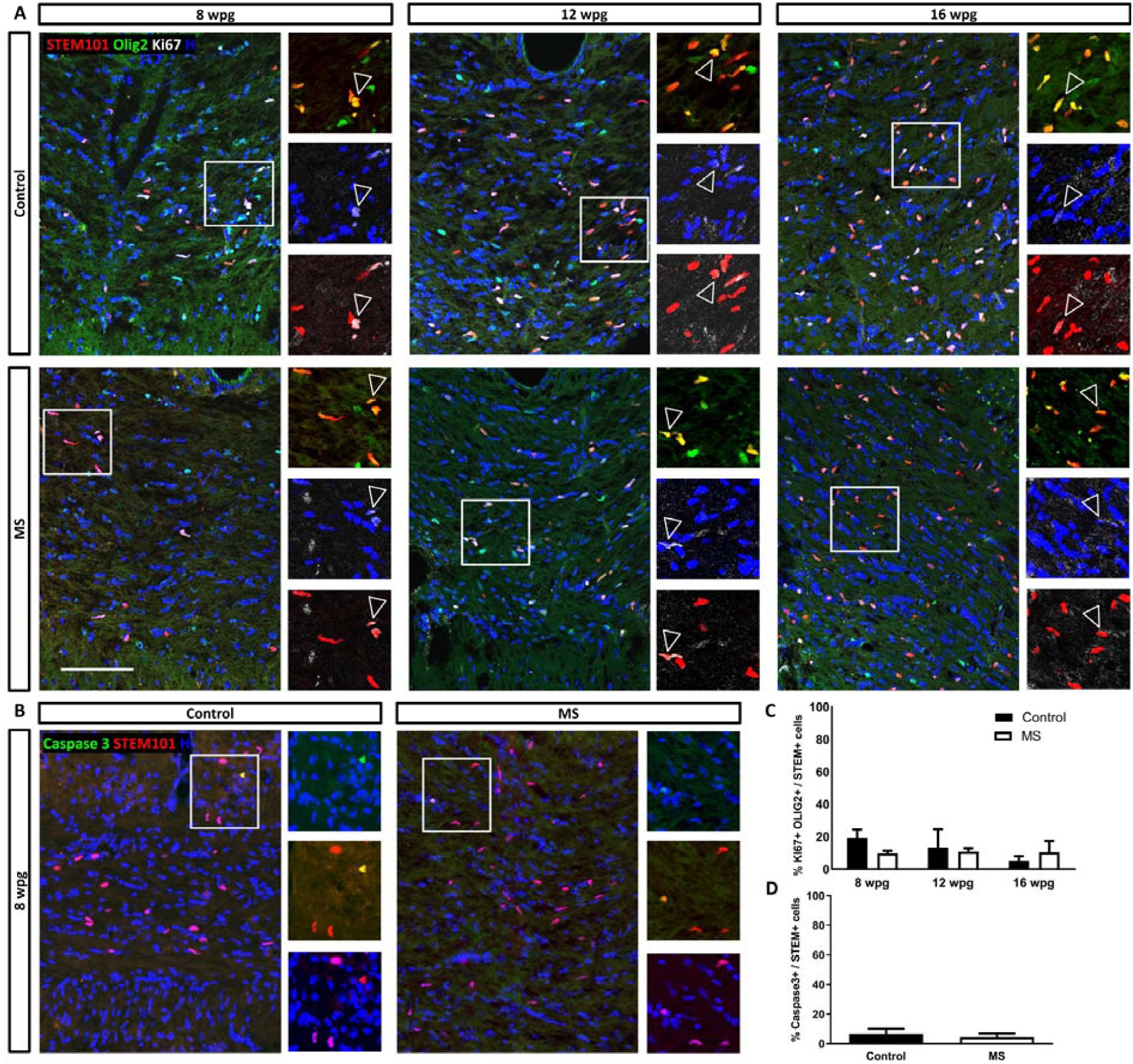
MS hiOL do not exhibit aberrant survival or proliferation in vivo. (**A** and **C**) Immunodetection of the human nuclei marker STEM101 (red) combined with OLIG2 (green) and the proliferation marker Ki67 (white), shows that a moderate proportion of MS-hiOLs sustains proliferation (empty arrowheads in the insets) following transplantation in their host developing brain, with no significant difference in the rate of proliferation between MS- and control-hiOLs over time. (**B** and **D**) Immunodetetion of the apoptotic marker Caspase3 (green) indicates that MS-hiOLs survive as well as control-hiOLs 8 wpg. Two-way ANOVA followed by Tukey’s multiple comparison or Mann-Withney t-tests were used for the statistical analysis (n=3–4 mice per group). Error bars represent SEMs. Wpg: weeks post graft. H, Hoechst dye. Scale bars: 100 μm.

### MS-hiOL do not show a differentiation block over time

Since MS-hiOLs proliferated and survived as well as control cells, we next questioned whether their differentiation potential into mature oligodendrocytes could be affected. We used the human nuclei marker STEM101 to detect all human cells in combination with SOX10, a general marker for the oligodendroglial lineage, and CC1 as a marker of differentiated oligodendrocytes. We found that the number of MS oligodendroglial cells (SOX10+) increased slightly but significantly with time, most likely resulting from sustained proliferation (Fig. 2A and B). Moreover, they timely differentiated into mature CC1+ oligodendrocytes with a 4-fold increase at 12 wpg, and a 5-fold increase at 16 wpg, when compared to 8 wpg, and with no difference with control-hiOLs (Fig. 2B and C).

**Fig. 2.**
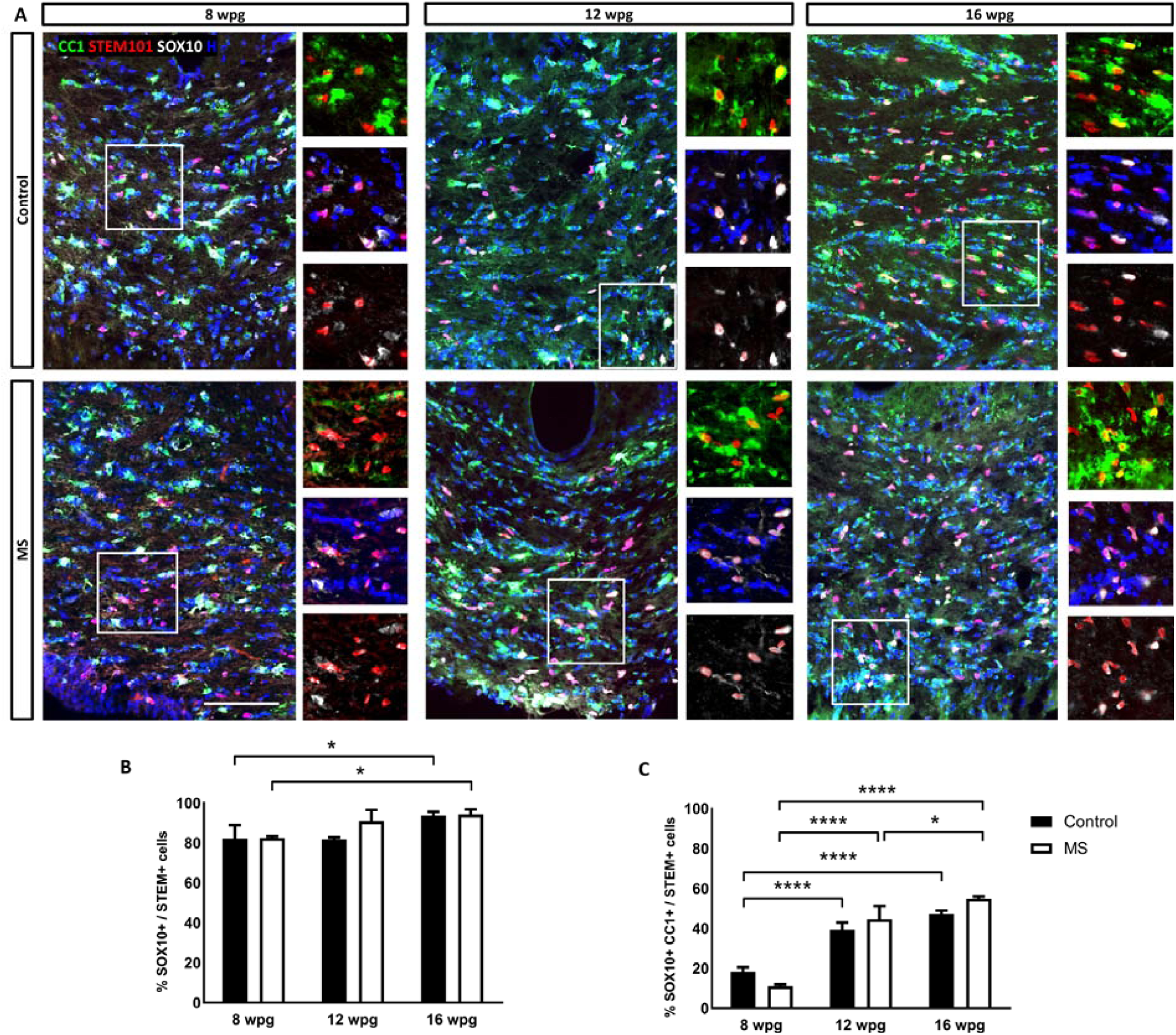
Differentiation of MS-hiOLs into mature oligodendrocytes is timely regulated in the corpus callosum of the developing Shi/Shi:Rag2-/- brain. (**A**) Combined immunodetection of human nuclei marker STEM101 (red) with CC1 (green) and SOX10 (white) for control (top) and MS-hiOLs (bottom) at 8, 12, and 16 wpg. (**B** and **C**) Quantification of SOX10+/STEM+ cells (**B**) and CC1+SOX10+ over STEM+ cells (**C**). While the percentage of human oligodendroglial cells increased only slightly with time, the percentage of mature oligodendrocytes was significantly time regulated for both MS- and control-hiOLs. Two-way ANOVA followed by Tukey’s multiple comparison tests were used for the statistical analysis of these experiments (n=3–4 mice per group). Error bars represent SEMs. *P < 0.05, and ****P < 0.0001. Wpg: weeks post graft, H: Hoechst dye; Scale bar: 100 μm.

### MS-hiOL do not show an aberrant pattern of myelination

The absence of abnormal MS-hiOL differentiation did not exclude a potential defect in myelination potential. We further investigated the capacity of MS-hiOLs to differentiate into myelin-forming cells. We focused our analysis on the core of the corpus callosum and fimbria. MS-hiOLs, identified by the human nuclear and cytoplasmic markers (STEM101 and STEM121), evolved from a bipolar to multi-branched phenotype (Fig. 3A and fig. S2: compare 4 wpg to 8 and 12 wpg), and differentiated progressively into MBP+ cells associated, or not, with T shaped MBP+ myelin-like profiles of increasing complexity (Fig. 3a, Fig. 4B, fig. S2). Myelin-like profiles clearly overlapped with NF200+ axons (Fig. 4A) and formed functional nodes of Ranvier expressing ankyrin G and flanked by paranodes enriched for CASPR (Fig. 4B) or neurofascin (Fig. 4C), as previously observed with control– hiOLs (*28*).

**Fig. 3.**
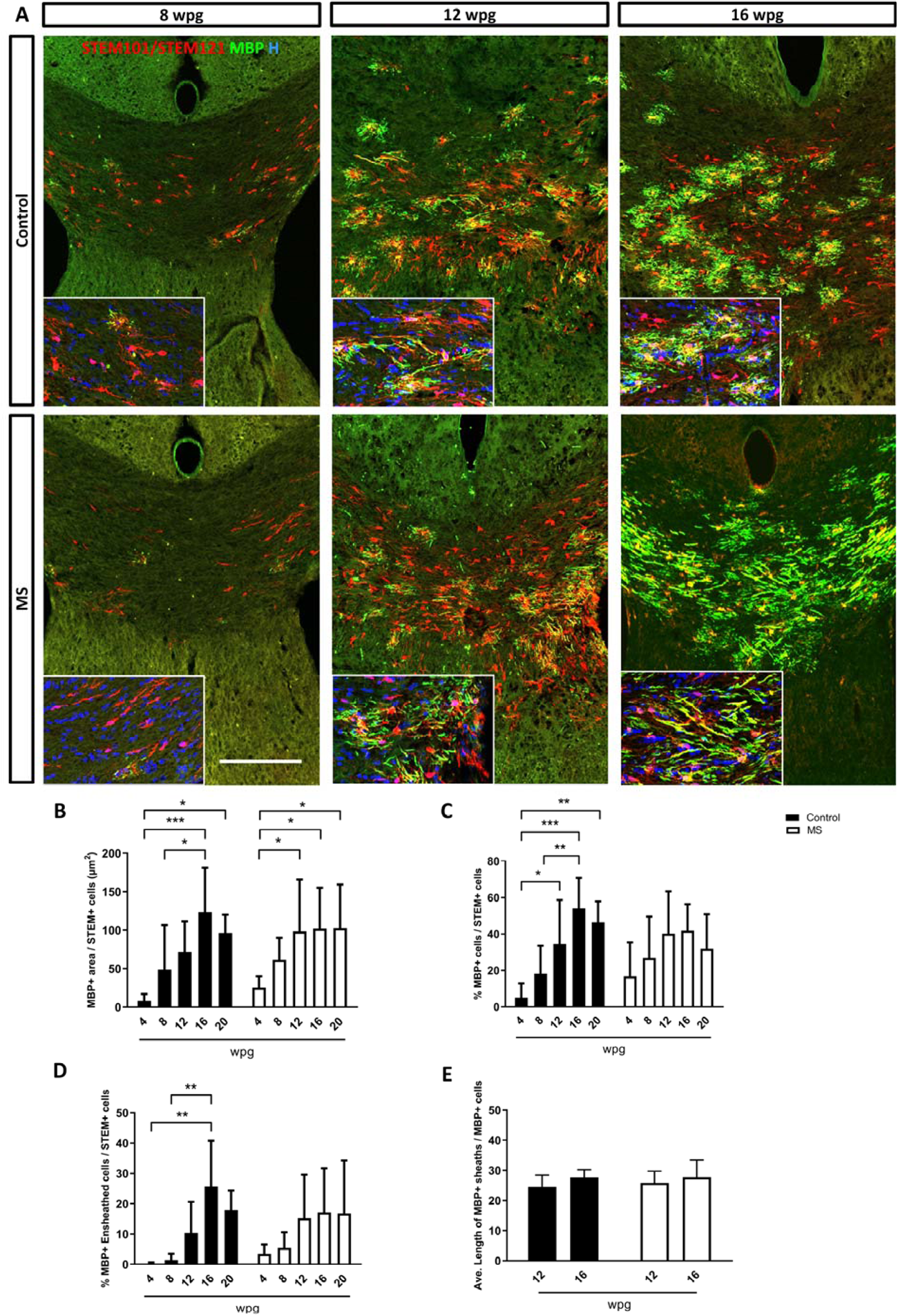
MS-hiOL derived progeny extensively myelinates the dysmyelinated Shi/Shi:Rag2-/- corpus callosum. (**A**) Combined detection of human nuclei (STEM101) and human cytoplasm (STEM 121) (red) with MBP (green) in the *Shi/Shi Rag2*^*-/-*^ corpus callosum at 8, 12 and 16 wpg. General views of horizontal sections at the level of the corpus callosum showing the progressive increase of donor derived myelin for control- (top) and MS- (bottom) hiOLs. (**B**) Evaluation of the MBP+ area over STEM+ cells. (**C** and **D**) Quantification of the percentage of (**C**) MBP+ cells and (**D**) MBP+ ensheathed cells. **e** Evaluation of the average sheath length (µm) per MBP+ cells. No obvious difference was observed between MS and control-hiOLs. Two-way ANOVA followed by Tukey’s multiple comparison tests were used for the statistical analysis of these experiments (n=6–14 mice per group). Error bars represent SEMs. *P < 0.05, **P < 0.01, and ***P < 0.001. Wpg: weeks post-graft. H, Hoechst dye. Scale bar: 200 μm. See also Figures S2 and S3.

**Fig. 4.**
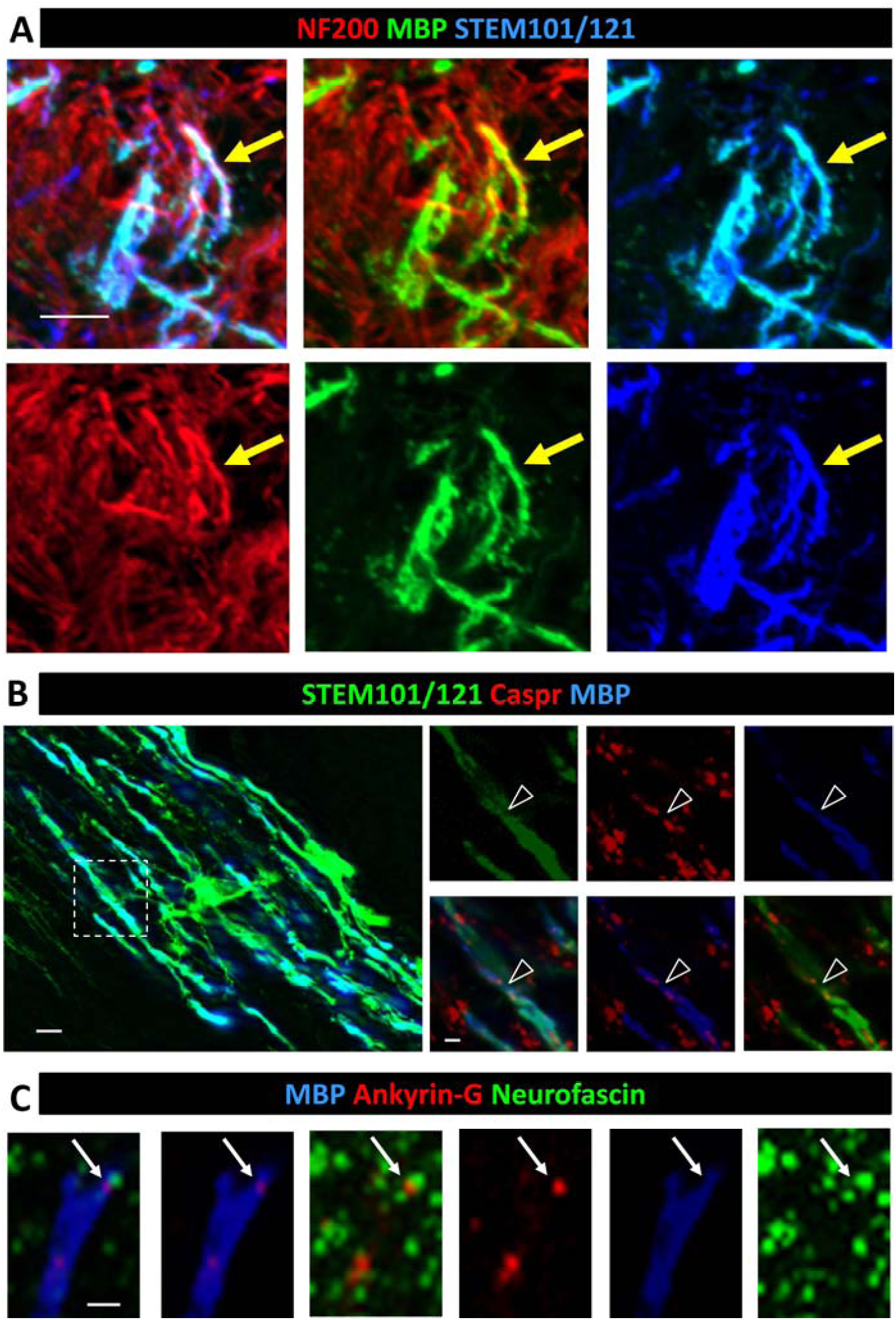
MS-hiOL derived myelin surrounds host axons and form bona-fide nodes of Ranvier. (**A**) Immunodetection of STEM101/121 (blue) with MBP (green) and Neurofilament NF200 (red) and identifies donor-derived myelin overlapping with NF200+ host axons 16 wpg. Arrows point to one example of co-labeling. (**B**) MS-derived oligodendrocytes (STEM101+/121+, green) connected to its T-shaped MBP+ myelin internodes (blue) flanked by paranodal Caspr+ domains (red). (**C**) MS-derived myelin (blue) integrated into axo-glial elements expressing nodal Ankyrin G (red) and paranodal neurofascin (green). Small arrows identify one example of node of Ranvier. Scale bars in **A** and **B**: 10 µm, in **C**: 2 µm.

We further analyzed, in depth, the myelinating potential of MS-hiOLs, applying automated imaging and analysis, providing multi-parametric quantification of individual hiOL in vitro (*32*) at 4, 8, 12, 16 and 20 wpg in vivo (Fig. 3B to D). We first examined the MBP+ surface area generated by the STEM+ cell population (Fig. 3B). While MS-hiOLs generated very low amount of myelin at 4 wpg, they generated significantly more myelin at 12, 16 and 20 wpg, with similar findings for control-hiOLs, highlighting the rapid progress in the percentage of myelin producing STEM+ cells in MS group over time.

We also quantified the percentage of STEM+ cells expressing MBP as well as the percentage of MBP+ with processes associated with linear myelin-like features, which we called MBP+ ensheathed cells. Both parameters increased significantly with time for control-hiOLs, reaching a plateau at 16wpg. The same tendency was achieved for MS-hiOLs with no significant differences between the control and MS-hiOL groups (Fig. 3C and D).

Myelin sheath length is considered to be an intrinsic property of oligodendrocytes (*33*). We analyzed this paradigm in our MS cohort at 12 and 16 wpg, time points at which sheaths were present at a density compatible with quantification. For those time-points, we found that the average MS MBP+ sheath length was equivalent to that of control with 25.86 ± 0.98 and 27.74 ± 1.52 µm for MS-hiOLs and 24.52 ± 1.48 and 27.65 ± 0.96 µm for control-hiOLs at 12 and 16 wpg, respectively (Fig. 3F). In summary our detailed analysis of immunohistochemically labeled sections indicates that, MS-hiOLs did not generate abnormal amounts of myelin in vivo, when compared to control-hiOLs.

Moreover, the myelinating potential of MS-hiOLs was further validated after engraftment in the developing spinal cord (4 weeks of age). Immunohistological analysis 12 wpg, revealed that STEM+ cells not only populated the whole dorsal and ventral columns of the spinal cord with preferential colonization of white matter, they also generated remarkable amounts of MBP+ myelin-like internodes that were found on multiple spinal cord coronal sections (fig. S3) thus indicating that their myelination potential was not restricted to only one CNS structure.

### MS derived myelin is compacted and rescues the slow trans-callosal conduction of their MBP deficient host

The presence of normal amounts of donor MBP+ myelin-like structures in the shiverer forebrain does not exclude potential structural anomalies. Therefore, we examined the quality of MS derived myelin at the ultrastructural level at 16 wpg in the Shi/Shi:Rag2-/- forebrain. In the corpus callosum of both MS and control-hiOLs grafted mice, we detected numerous axons surrounded by electron dense myelin, which at higher magnification was fully compacted compared to the uncompacted shiverer myelin (Fig. 5A to F); (*28, 34*). Moreover, MS myelin reached a mean g-ratio of 0.76 ± 1.15 comparable to that of control myelin (0.75 ± 1.56) (Fig. 5G) and thus a similar myelin thickness. This argues in favor of 1) MS-hiOLs having the ability to produce normal compact myelin and thus its functional normality and 2) a similar rate of myelination between the two groups, and consequently, an absence of delay in myelination for MS-hiOLs.

**Fig. 5.**
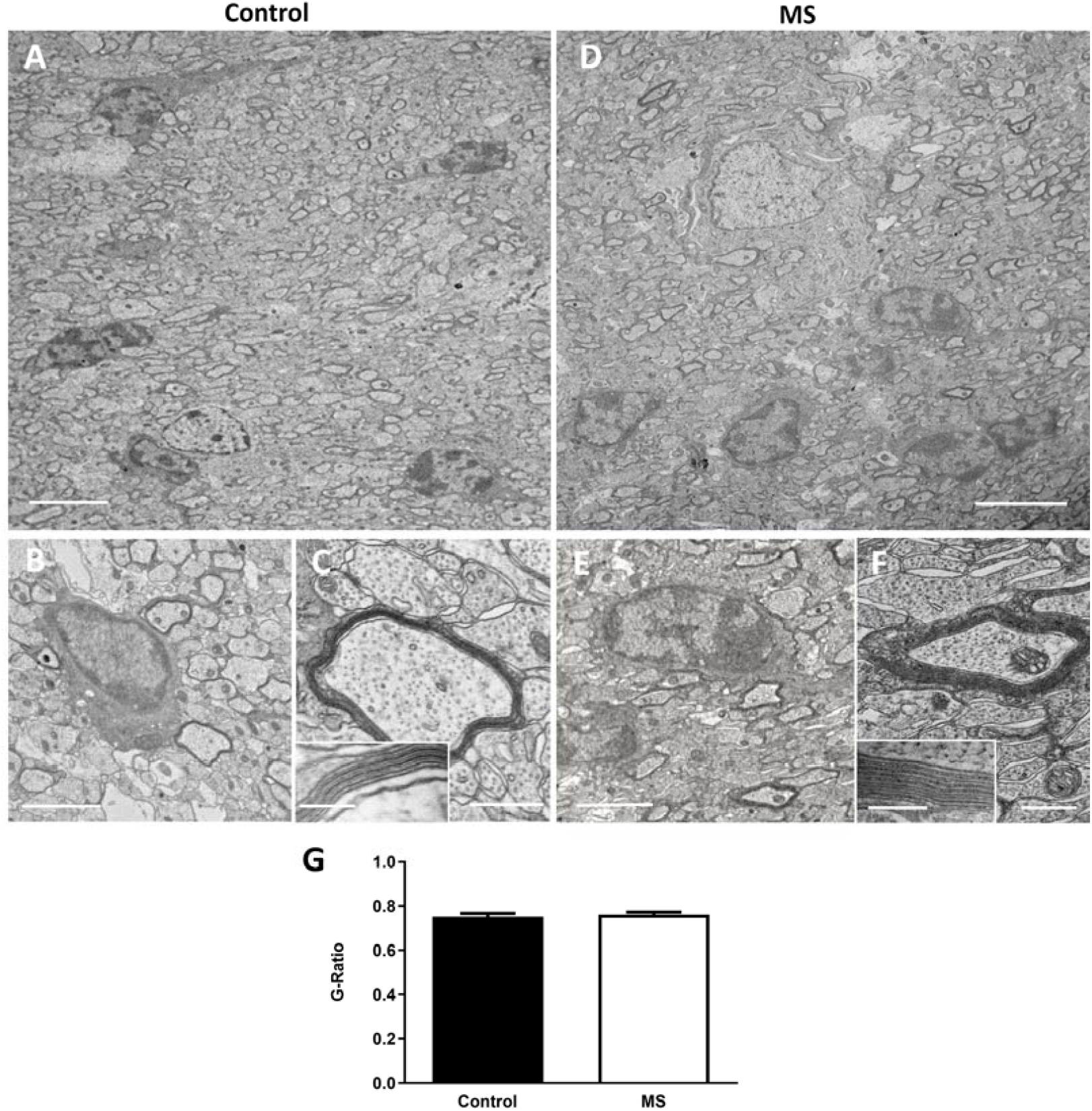
MS-hiOLs produce compact myelin in the dysmyelinated Shi/Shi:Rag2 -/- corpus callosum. (**A** to **F**) Ultrastructure of myelin in sagittal sections of the core of the corpus callosum 16 wpg with control-hiOLs (**A** to **C**) and MS-hiOLs (**D** to **F**). (**A** and **D**) General views illustrating the presence of some electron dense myelin, which could be donor-derived. (**B, C, E** and **F**) Higher magnifications of control (**B** and **C**) and MS (**E** and **F**) grafted corpus callosum, validate that host axons are surrounded by thick and compact donor derived myelin. Insets in **C**, and **F** are enlargements of myelin, and show the presence of the major dense line. No difference in compaction and structure is observed between the MS and control myelin. (**G**) Quantification of g-ratio revealed no significant difference between myelin thickness of axons myelinated by control and MS-hiOLs. Mann-Withney t-tests was used for the statistical analysis of this experiment (*n* = 4 mice per group). Error bars represent SEMs. Scale bars in **A** and **D**: 5 μm, in **B** and **E**: 2 µm, in **C** and **F**: 500 nm (with 200 nm and 100 nm respectively in **C** and **F** insets).

Myelin compaction has a direct impact on axonal conduction with slower conduction in shiverer mice compared to wild-type mice (*12, 35*). We therefore questioned whether newly formed MS-hiOLs derived myelin has the ability to rescue the slow axon conduction velocity of shiverer mice in vivo (Fig. 6). As previously performed with fetal glial restricted progenitors (*13*), trans-callosal conduction was recorded in vivo, at 16 wpg in mice grafted with MS- and control-hiOLs, and compared with non-grafted shiverer and wild-type mice. As expected, conduction in non-grafted shiverer mice was significantly slower compared to wild-type mice. However, axon conduction velocity was rescued by MS-hiOLs, and to the same extent as by control-hiOLs.

**Fig. 6.**
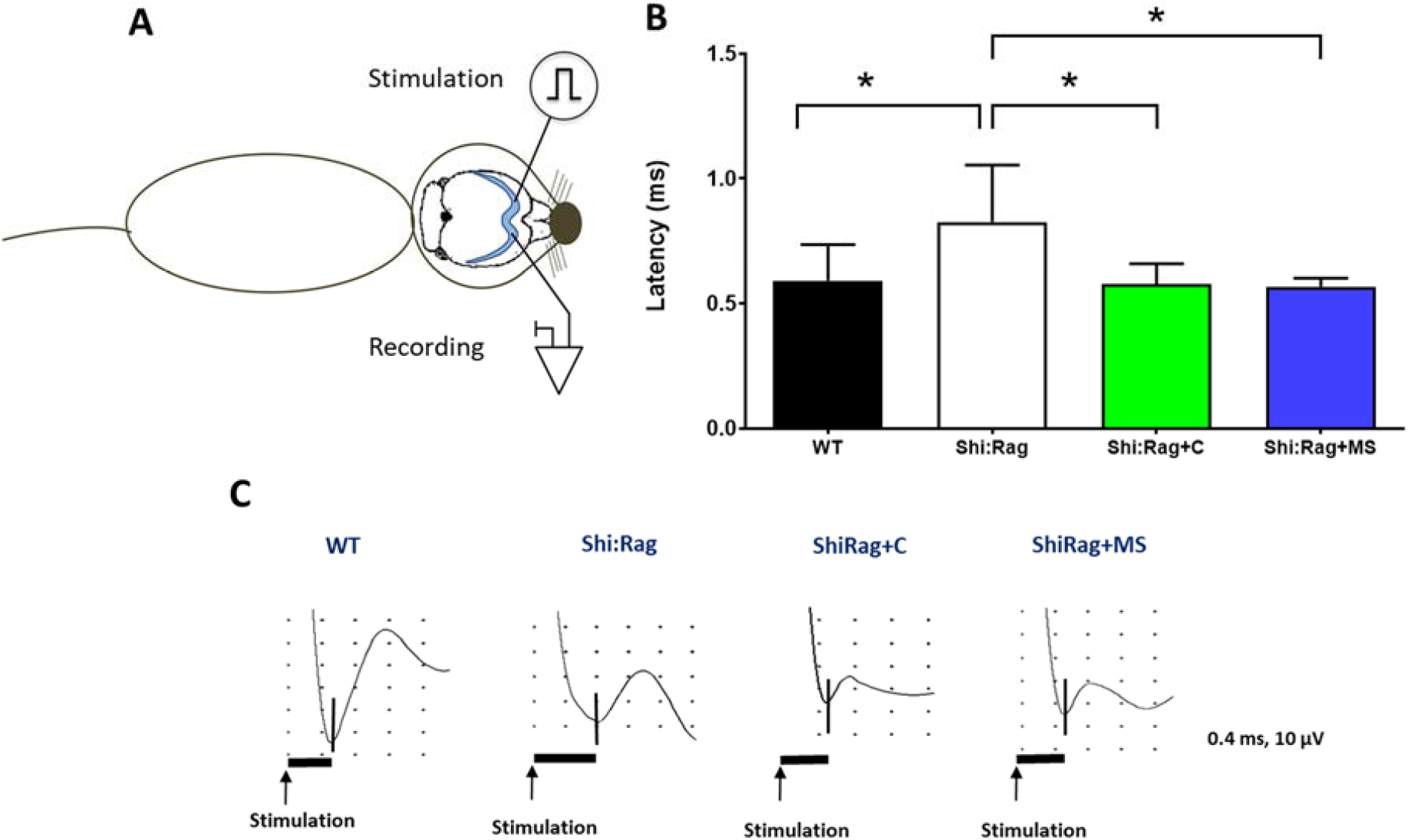
MS-hiOLs improve trans-callosal conduction of the dysfunctional Shi/Shi:Rag2-/- axons to the same extent than control-hiOLs. (**A**) Scheme illustrating that intra-callosal stimulation and recording are performed in the ipsi- and contra-lateral hemisphere respectively. (**B**) N1 latency was measured following stimulation in different groups of *Shi/Shi:Rag2*^*-/-*^ : intact or grafted with control or MS-hiOLs and wild-type (WT) mice, at 16 wpg. MS-hiOL derived myelin significantly restored trans-callosal conduction latency in *Shi/Shi:Rag2*^*-/-*^ mice to the same extent than control-derived myelin (*P* = 0.01) and close to that of WT levels. One-way ANOVA with Dunnett’s multiple comparison test for each group against the group of intact *Shi/Shi:Rag2*^*-/-*^ was used. Error bars represent SEMs. *P < 0.05. (**C**) Representative response profiles for each group. Scales in Y axis is equal to 10 µV and in the X axis is 0.4 ms.

### Grafted MS-hiOL show typical cell stage-specific electrophysiological properties

Rodent oligodendrocyte progenitors and oligodendrocytes can be distinguished by cell stage-specific electrophysiological properties (*36, 37*). To assess the electrophysiological properties of oligodendroglial lineage cells derived from human grafted control and MS-hiOLs, RFP-hiOLs were engrafted in the Shi/Shi:Rag2-/-forebrain, and recorded with a K-gluconate-based intracellular solution in acute corpus callosum slices at 12-15 wpg (Fig. 7A). As previously described for rodent cells, hiOLs in both groups were identified by their characteristic voltage-dependent current profile recognized by the presence of inward Na+ currents and outwardly rectifying steady-state currents (Fig. 7B). We found that ∼60% and ∼44% of recorded cells were oligodendrocyte progenitors derived from MS and control progenies, respectively. No significant differences were observed in the amplitude of Na+ currents measured at -20 mV (Fig. 7D) or steady-state currents measured at +20 mV between MS- and control-derived oligodendrocyte progenitors (Isteady=236.70±19.45 pA and 262.10±31.14 pA, respectively; p=0.8148, Mann Whitney U test). We further confirmed the identity of these cells by the combined expression of SOX10 or OLIG2 with STEM101/121, and the absence of CC1 in biocytin-loaded cells (Fig. 7F, top). The remaining recorded cells (MS and control) did not show detectable Na+ currents after leak subtraction and were considered to be differentiated oligodendrocytes by their combined expression of SOX10, STEM101/121 and CC1 in biocytin-loaded cells (Fig. 7F, bottom). The I-V curve of these differentiated oligodendrocytes displayed a variable profile that gradually changed from voltage-dependent to linear as described for young and mature oligodendroglial cells in the mouse (*36*). Figure 7C illustrates a typical linear I-V curve of a fully mature MS-derived oligodendrocytes. No significant differences were observed in the amplitude of steady-state currents measured at +20 mV between MS- and control-derived oligodendrocytes (Fig. 7E).

**Fig. 7.**
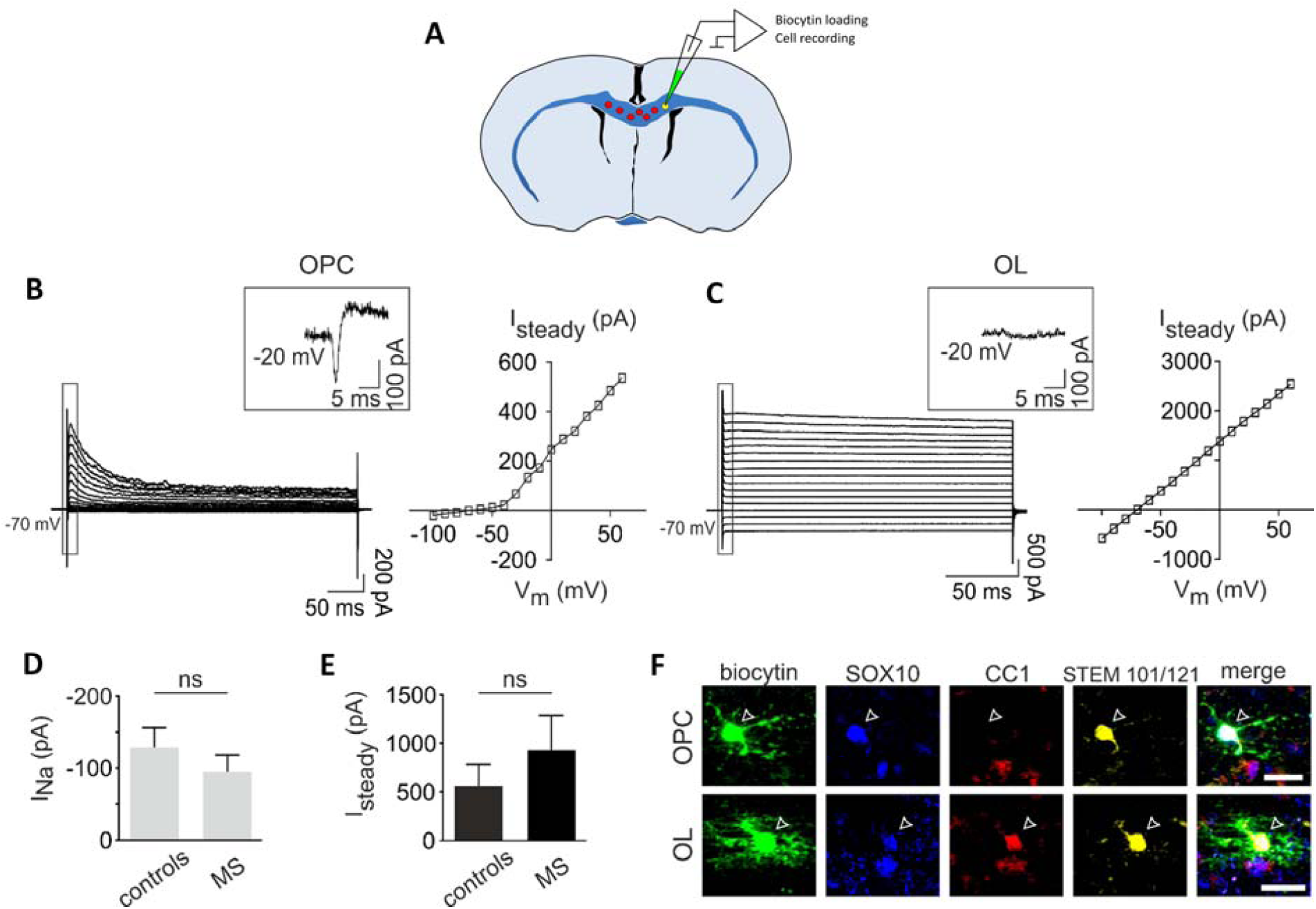
Grafted MS-hiOL show typical cell stage-specific electrophysiological properties. (**A**) Schematic representation of the concomitant Biocytin loading and recording of single RFP+ hiOL derivative in an acute coronal brain slice prepared from mice engrafted with hiOLs (control or MS) and analyzed at 12-14wpg. (**B** and **C**) Currents elicited by voltage steps from -100 mV to +60 mV in a control-oligodendrocyte progenitor (**B**, left) and a MS-oligodendrocyte (**C**, left). Note the presence of an inward Na^+^ current obtained after leak subtraction in the oligodendrocyte progenitor, but not in the oligodendrocyte (insets). The steady-state I-V curve of the oligodendrocyte progenitor displays an outward rectification (**B**, right) while the curve of the oligodendrocyte has a linear shape (**c**, right). (**D**) Mean amplitudes of Na^+^ currents measured at -20 mV in control and MS iPSCs-derived oligodendrocyte progenitors (n=8 and n=9, respectively; p=0.743, Mann-Whitney U test). (**E**) Mean amplitudes of steady-state currents measured at +20 mV in control and patient differentiated iPSCs-derived oligodendrocytes (n=10 and n=6, respectively; p=0.6058, Mann-Whitney U test). (**F**) A control iPSCs-derived oligodendrocyte progenitor loaded with biocytin and expressing OLIG2, STEM101/121 and lacking CC1 (top panels) and a MS iPSC derived oligodendrocyte loaded with biocytin and expressing SOX10, CC1, STEM101/121 (bottom panels). Scale bar: 20 µm.

Overall, the electrophysiological profile of oligodendrocyte progenitors and oligodendrocytes derived from control and MS were equivalent and showed similar characteristics to murine cells (*36, 37*).

### MS- and control-hiOL progeny functionally connect to other glial cells

Studies with rodents have reported that oligodendrocytes exhibit extensive gap-junctional intercellular coupling between other oligodendrocytes and astrocytes (*38, 39*). Whether oligodendrocytes derived from grafted human cells can be interconnected with cells in the adult host mouse brain was not known, and whether MS-hiOLs maintain this intrinsic property was also not addressed. Since biocytin can pass through gap junctions, we inspected biocytin-labeled cells for dye coupling Fig. 7A, Fig. 8A and B).

**Fig. 8.**
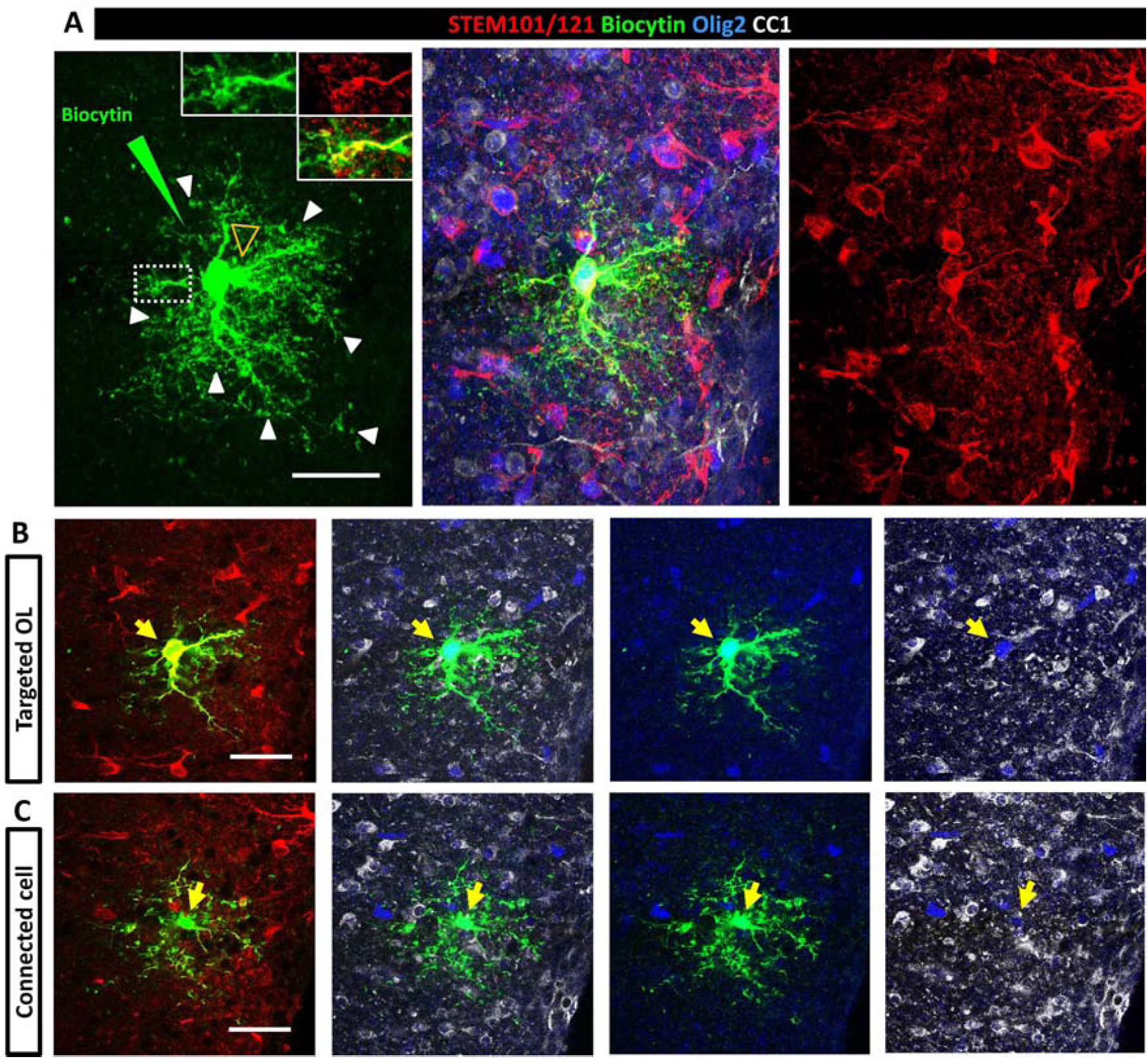
Grafted hiOL derivatives are functionally connected to murine cells. (**A**) Z-stack identifying a target and connected cell. One given single grafted human RFP+ cell (per acute slice) was loaded with biocytin by a patch pipette and allowed to rest for 30 min. The white arrowheads and insets in **A** illustrate biocytin diffusion up to the donut-shaped tip of the human oligodendrocyte processes. Another biocytin-labeled cell (empty yellow arrowhead) was revealed at different morphological level indicating diffusion to a neighboring cell and communication between the two cells via gap junctions. (**B** and **C**) Split images of **a** showing the target (**B**) and connected (**C**) cell separately at different levels. Immunolabeling for the combined detection of the human markers STEM101/121 (red), OLIG2 (blue) and CC1 (white) indicated that the target cell is of human origin (STEM+) and strongly positive for OLIG2 and CC1, a mature oligodendrocyte, and that the connected cell is of murine origin (STEM-) and weakly positive for OLIG2 and CC1, most-likely an immature oligodendrocyte. Scale bars: 30 µm. See also Fig. 9.

We found that 2 out of 7 MS-derived oligodendrocytes (∼29%) and 5 out of 21 control-derived oligodendrocytes (∼24%) were connected with a single neighboring cell, which was either human or murine (Fig. 8), except in one case where three mouse cells were connected to the biocytin-loaded human cell. These findings reveal that gap junctional coupling can occur between cells from the same or different species, and MS-hiOLs can functionally connect to other glial cells to the same extent as their control counterparts.

To validate the presence of glial-glial interactions, we investigated whether the grafted hiOL-derived progeny had the machinery to be connected to one another via gap junctions. To this end, we focused on oligodendrocyte-specific Cx47 and astrocyte-specific Cx43 as Cx43/47 channels, which are important for astrocyte/oligodendrocyte cross talk during myelination and demyelination (*40, 41*). Combined immunolabeling for hNOGOA, CC1, OLIG2 and Cx47 revealed that MS-derived oligodendrocyte cell bodies and processes were decorated by Cx47+ gap junction plaques, that were often shared by exogenous MS-derived oligodendrocytes or by MS and endogenous murine oligodendrocytes (Fig. 9A). In addition, co-labeling exogenous myelin for MBP and Cx43 identified the presence of several astrocyte-specific CX43 gap junction plaques between human myelin internodes, highlighting contact points between astrocyte processes and axons at the human-murine chimeric nodes of Ranvier (Fig. 9B).

**Fig. 9.**
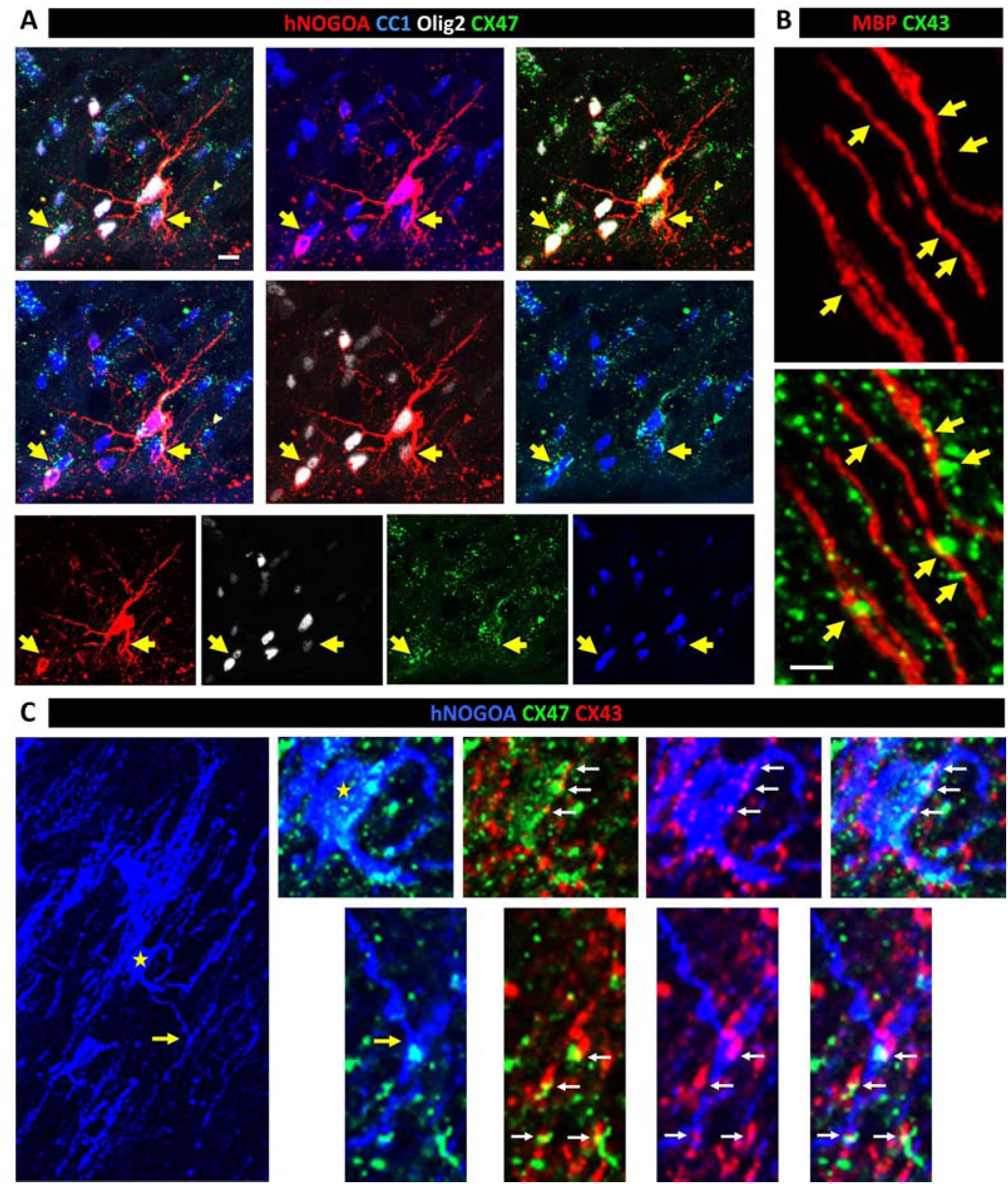
Grafted MS-hiOL derivatives express the gap junction plaques Cx47, which are associated with the astrocyte-gap junction plaque Cx43. (**A**) Immunodetection of hNOGOA (red), CC1 (blue), OLIG2 (white) and oligodendrocyte Cx47, illustrates the presence of a hiOL-derived mature oligodendrocyte, which processes are decorated by Cx47 gap junction plaques. Yellow arrows point to examples of plaques between two adjacent oligodendrocytes with human-host origins. (**B**) Combined immunodetection of human MBP+ myelin and astrocyte specific Cx43, shows Cx43 expression where MBP+ internodes are interrupted (yellow arrows), presumably at nodes of Ranvier. (**C**) Illustrates a hNOGOA+ oligodendrocyte (blue) expressing oligodendrocyte-specific Cx47+ gap junction plaques (green) juxtaposed to astrocyte-specific Cx43+ gap junction plaques (red), suggesting human oligodendrocyte-murine astrocyte connections at the levels of human oligodendrocyte body (yellow star) or at the T-shape myelin-like structure (yellow arrow) of the hNOGOA+ oligodendrocyte (blue). White arrows point to example of Cx47-Cx43 associations. Scale bars: 10 µm for **A**, 5 µm for **B**, 20 µm for **C**.

Finally, co-labeling of hNOGOA, with CX47 and the astrocyte-specific Cx43, revealed co-expression of oligodendrocyte- and astrocyte-specific connexins at the surface of MS-derived oligodendrocyte cell bodies as well as at the level of T-shaped myelin-like structures (Fig. 9C), thus implicating connections between human oligodendrocytes and murine and/or human astrocytes as a small proportion of the grafted hiOLs differentiated into astrocytes. Immunolabeling for human anti-GFAP, and Cx43 showed that these human astrocytes were decorated by Cx43+ aggregates, as observed in the host subventricular zone (fig. S4A).

Furthermore, immunolabeling for human GFAP, mouse GFAP and Cx43 indicated that Cx43+ gap junctions were shared between human and mouse astrocytes as observed at the level of blood vessels (fig. S4B). These data validate interconnections between the grafted-derived human glia (MS and controls) with murine host glial cells, and confirm their interconnection with the pan-glial network.

## Discussion

Two main hypotheses have been long considered in understanding MS pathology and etiology: the outside-in hypothesis highlighting the role of immune regulators as extrinsic key players in MS pathology and possibly its repair failure, or the inside-out hypothesis pointing to the intrinsic characteristics of neuroglia including oligodendroglial cells as the main contributors in the MS scenario. Single-cell transcriptomic analysis revealed the presence of disease-specific oligodendroglia expressing susceptibility genes in MS brains (*18*), and altered oligodendroglia heterogeneity in MS (*19*). The question remains open as to whether these altered oligodendroglial phenotypes are acquired in response to the disease environment, or whether they reflect intrinsic traits of the MS oligodendroglial population. On the other hand, the whole exome sequencing analysis in 132 patients from 34 multi-incident families identified 12 candidate genes of the innate immune system and provided the molecular and biological rational for the chronic inflammation, demyelination and neurodegeneration observed in MS patients (*42*).

While none of these hypotheses have been fully proven or rejected, research efforts for a better understanding of this multifactorial disease have continued. Impaired remyelination or oligodendrocyte differentiation block in MS is still considered a potentially disease-relevant phenotype (*43, 44*). Many histological and experimental studies suggest that impaired oligodendrocyte progenitor to oligodendrocyte differentiation may contribute to limited remyelination in MS, although this view is challenged by recent studies questioning the contribution of oligodendroglial cells to remyelination (*19, 45, 46*). Understanding MS oligodendrocyte biology has been challenging mainly due to the following reasons: 1) oligodendroglial cells are not easily accessible to be studied in vivo; 2) dynamic remyelination observed in MS patients - that points to their individual remyelination potential - is inversely correlated with their clinical disability (*5*) highlighting even more complexity in oligodendrocyte heterogeneity between MS patients; 3) exclusion of the role of immune system players in understanding MS oligodendrocyte biology being inevitable in most of clinical or experimental studies.

In such a multifactorial disease, one of the most accessible and applicable approaches to overcome these problems is the generation of large quantities of bona-fide disease and control oligodendroglia using iPSC technology, and to investigate their genuine behavior in vivo after engraftment in a B and T cell-free system. One might consider that pluripotency induction could by in vitro manipulation, erase cell epigenic traits, and as a result, modulate their characteristics. While we cannot prove or reject this possibility at the present time, many reports have highlighted the different behavior of diseased iPSC-derived oligodendrocytes in comparison to those from healthy controls using the same technology in multifactorial diseases such as Schizophrenia (*21, 22*), Huntington’s disease (*20*) and others (*47*). In this regard, direct reprogramming of somatic cells into the desired cell type, bypassing the pluripotent stage, could be an attractive alternative. However, so far only mouse fibroblasts have been successfully directly converted into oligodendroglial cells, and with relatively low efficiency (*48, 49*).

In this study using a very efficient reprogramming method (*28*), and a purely dysmyelinating Shi/Shi:Rag2-/- mouse model to avoid confounding immune-mediated extrinsic effect, we show that MS-hiOLs derivatives survive, proliferate, migrate and timely differentiate into bona-fide myelinating oligodendrocytes in vivo as efficiently as their control counterparts. Nicaise and colleagues reported that iPSC-NPCs from PPMS cases did not provide neuroprotection against active CNS demyelination compared to control iPSC-NPCs lines (*23*), and failed to promote oligodendrocyte progenitor maturation due to senescence without affecting their endogenous capacity to generate myelin-forming oligodendrocytes (*24, 25*). However, their myelinating potential was not evaluated against control cells. Generation of iPSC-oligodendrocyte progenitors from PPMS or RRMS patients has also been reported by other groups, yet with no evidence for their capacity to become functional oligodendrocytes in vivo (*26, 27*). Thus, so far, no conclusion can be made regarding the potential impact of disease severity (PPMS verses RRMS) on the functionality of the iPSC-derived progeny.

Here, we compared side-by-side, and at different time points after engraftment, hiOLs from RRMS patients and controls including two pairs of homozygous twins discordant for disease and found no significant difference in their capacity to timely differentiate (according to the human tempo of differentiation) and efficiently myelinate axons in the shiverer mouse in terms of the percentage of MBP+ cells generated, amount of myelin produced, length of MBP+ sheaths, as well as the ultrastructure and thickness of myelin sheaths. MS-hiOLs also re-constructed nodes of Ranvier expressing nodal components key to their function. We not only verified that the grafted MS-hiOLs derivatives were anatomically competent, but also established their functionality at the electrophysiological level using 1) in vivo recordings of transcallosal evoked potentials, and 2) ex-vivo recordings of the elicited current-voltage curves of the grafted MS-hiOLs verses controls. Our data show that the grafted MS-hiOLs were able to rescue the established delayed latency in the shiverer mice to the same extent as control cells, as previously reported for human fetal glial progenitors grafted in the same model (*13*). Moreover, at the single cell level, MS-hiOL-derived oligodendrocyte progenitor and oligodendrocytes did not harbor aberrant characteristics in membrane currents as compared to those of control cells ex vivo. Thus, iPSC-derived human oligodendroglial cells shift their membrane properties with maturation as previously observed in vitro (*50*) and these properties are not impaired in MS.

While most pre-clinical transplantation studies have focused on myelination potential as the successful outcome of axo-glia interactions, less is known about the capacity of the grafted cells to fulfill glial-glial interactions in the pan-glial syncytium, which could ensure maintenance of newly generated myelin (*51*) and cell homeostasis (*52, 53*). Oligodendrocytes are extensively coupled to other oligodendrocytes, and oligodendrocyte progenitors through the homologous gap junctions Cx47 (*38*). These intercellular interactions between competing oligodendroglial cells influence the number and length of myelin internodes as well as the initiation of differentiation (*54, 55*). Oligodendrocytes are also coupled to astrocytes through heterologous gap junctions such as Cx32/Cx30 and Cx47/Cx43 (*56*). Disruption of oligodendrocytes from each other and from astrocytes, i.e. deconstruction of pan-glial network, has been observed in experimental models of demyelination (unpublished data) and frequently reported in MS and neuromyelitis optica (*41, 57-59*). Mutations in Cx47 and Cx32 result in developmental CNS and PNS abnormalities in leukodystrophies (*60, 61*). Moreover, experimental ablation of Cx47 results in aberrant myelination (*62*) and significantly abolished coupling of oligodendrocytes to astrocytes (*38*).

In view of the major role of Cx-mediated gap junctions among oligodendrocytes and between oligodendrocytes and astrocytes during myelin formation (*56*), we asked whether MS-hiOL progeny was capable of making functional gap junctions with other glial cells, and integrating into the host panglial network. We show that grafted MS-hiOLs, in common with rodent oligodendrocytes, express Cx47 that was frequently shared between the human and murine oligodendrocytes (through Cx47-Cx47), but also in conjunction with the astrocyte Cx43 (via Cx47/Cx43). The dye-coupling study highlighted that MS-hiOLs, similarly to control cells, were capable of forming functional gap junctions with neighbor murine or human glial cells, indicating that MS-hiOLs retained the intrinsic property, not only to myelinate host axons, but also to functionally integrate into the host pan-glial network. While our study focused mainly on oligodendroglial cells, a small proportion of the grafted hiOLs differentiated into astrocytes expressing Cx43. Interestingly, these human astrocytes were detected associated with blood vessels, or the subventricular zone, where they were structurally gap-junction-coupled to mouse astrocytes as previously observed after engraftment of human fetal glial restricted progenitors (*63*).

Altogether our data highlight that human skin derived glia retain characteristics of embryonic/fetal brain-derived glia as observed for rodent cells (*12*). In particular, we show that MS-hiOLs timely differentiate into mature oligodendrocytes, functionally myelinate host axons and contribute to the human-mouse chimeric pan-glial network as efficiently as control-hiOLs. These observations favor a role for extrinsic rather than intrinsic oligodendroglial factors, in the failed remyelination of MS. Interestingly, the international Multiple Sclerosis genetics consortium after analyzing the genomic map of more than 47000 MS cases and 63000 control subjects, implicated microglia as well as multiple different peripheral immune cell populations in disease onset (*64*). Moreover, neuroinflammation appears to block oligodendrocyte differentiation and to alter their properties to aggravate the auto-immune process (*65*). Further, MS lymphocytes are reported to exhibit intrinsic capacities that drive myelin repair in a mouse model of demyelination (*66*). On the other hand, recent data highlighted the presence of disease-specific oligodendroglia in MS (*18, 19*), However, it should be considered that most of the data were collected using single nuclei RNA-sequencing of postmortem tissues from MS or control subjects of different ages that were suffering from other disorders ranging from cancer to sepsis and undergoing various treatment - and so died for different reasons - that may have influenced the type or level of RNA expression by the cells.

Data from the present study provide valuable findings in the scenario of MS pathology highlighting that MS-hiOLs, regardless of major manipulators of the immune system, do not lose their intrinsic capacity to functionally myelinate and interact with other neuroglial cells in the CNS under non-pathological conditions. Whether MS-hiOLs or oligodendroglial cells directly reprogrammed from MS fibroblasts would behave similarly well, if challenged with neuropathological inflammatory conditions as opposed to conditions wherein the immune system is intact (presence of T-cell and B-cells), will require further investigation.

In sum, the findings obtained provide valuable insights not only into the biology of MS oligodendroglia, but also their application for cell-based therapy, and should contribute to the establishment of improved preclinical models for in vivo drug-screening of pharmacological compounds targeting the OPCs, OLs and their interactions with the neuronal and pan-glial networks.

## Materials and Methods

### Study design

We examined side-by-side the molecular, cellular and functional behavior of MS hiOLs with their control counterparts after their engraftment in a dysmyelinating animal model to avoid the effect of major immune modulators. We used 3 MS and 3 control cell lines including two monozygous twin pairs discordant for the disease. We performed in vivo studies in mouse with sample size between three to six per cell line/time point/assay required to achieve significant differences. Numbers of replicates are listed in each figure legend. Animals were monitored carefully during all the study time and animal welfare criteria for experimentation were fully respected. All experiments were randomized with regard to animal enrollment into treatment groups. The same experimenter handled the animals and performed the engraftment experiments to avoid errors. The data were analyzed by a group of authors.

### Animals

Shiverer mice were crossed to Rag2 null immunodeficient mice to generate a line of Shi/Shi:Rag2-/- dysmyelinating-immunodeficient mice in order to 1) prevent rejection of the grafted human cells and allow detection of donor-derived wild-type myelin and 2) investigate the original behavior of MS-derived oligodendrocytes in a B cell/T cell-free environment. Mice were housed under standard conditions of 12-hour light/ 12-hour dark cycles with ad libitum access to dry food and water at the ICM animal facility. Experiments were performed according to European Community regulations and INSERM ethical committee (authorization 75-348; 20/04/2005) and were approved by the local Darwin ethical committee.

### Human cells

Fibroblasts were obtained under informed consent from 3 control and 3 RRMS subjects including two monozygous twin pairs discordant for the disease. They were reprogrammed into iPSCs using the replication incompetent Sendaï virus kit (Invitrogen) according to manufacturer’s instructions. Table S1 summarizes information about the human cell lines used in this study. The study was approved by the local ethical committees of Münster and Milan (AZ 2018-040-f-S, and Banca INSpe).

Human iPSCs were differentiated into neural precursors (NPC) by treatment with small molecules as described (*67, 68*). Differentiation of NPCs into O4+ oligodendroglial cells utilized a poly-cistronic lentiviral vector containing the coding regions of the human transcription factors Sox10, Olig2 and Nkx6.2 (SON) followed by an IRES-pac cassette allowing puromycin selection for 16h (*28*). For single cell electrophysiological recordings, the IRES-pac cassette was replaced by a sequence encoding red fluorescein protein (RFP). Briefly, human NPCs were seeded at 1.5 x 105 cells/well in 12-well plates, allowed to attach overnight and transduced with SON lentiviral particles and 5 µg/ml protamine sulfate in fresh NPC medium. After extensive washing, viral medium was replaced with glial induction medium (GIM). After 4 days, GIM was replaced by differentiation medium (DM). After 12 days of differentiation, cells were dissociated by accutase treatment for 10 min at 37°C, washed with PBS, re-suspended in PBS/0.5% BSA buffer and singularized cells filtered through a 70 µm cell strainer (BD Falcon). Cells were incubated with mouse IgM anti-O4-APC antibody (Miltenyi Biotech) following the manufacturer’s protocol, washed, re-suspended in PBS/0.5% BSA buffer (5 x 106 cells/ml) and immediately sorted using a FACS Aria cell sorter (BD Biosciences). Subsequently, human O4+ hiOLs were frozen and stored at -80°C. Media details were provided in (*28*).

### Cell transplantation

To assay hiOL contribution to forebrain developmental myelination, newborn Shi/Shi:Rag2-/- pups (n=148) were cryo-anesthetized and control and RRMS hiOLs transplanted bilaterally, rostral to the corpus callosum. Injections (1µl in each hemisphere, 105 cells/µl) were performed 1 mm caudally, 1 mm laterally from the bregma and to a depth of 1 mm as previously described (*69, 70*). Animals were sacrificed at 4, 8, 12, 16 and when indicated 20 wpg for immunohistological studies, and at one time-point for electron microscopy (16 wpg), ex vivo (12-15 wpg) and in vivo (16 wpg) electrophysiology.

To assay the fate of hiOLs in the developing spinal cord, 4 week old mice (n=4) were anaesthetized by intraperitoneal injection of a mixture of 100 mg/kg Ketamine (Alcyon) and 10 mg/kg Xylazine (Alcyon) and received a single injection at low speed (1µl/2min) of hiOLs (1 µl, 105 cells/μl) at the spinal cord thoracic level using a stereotaxic frame equipped with a micromanipulator and a Hamilton syringe. Animals were sacrificed at 12 wpg for immunohistological studies.

### Post mortem analysis

#### Immunohistochemistry

Shi/Shi:Rag2-/- mice grafted with control and RRMS hiOLs (n=3-6 per group and time-point) were sacrificed by trans-cardiac perfusion-fixation with 4% PFA in PBS. Tissues were post-fixed in the same fixative for 1 hr, and incubated in 20% sucrose in 1xPBS overnight before freezing at -80°C. Serial horizontal brain and spinal cord cross sections of 12 µm thickness were performed with a cryostat (CM3050S; Leica). Transplanted hiOLs were identified using anti-human cytoplasm (STEM121; Takara, Y40410, IgG1, 1:100), anti-human nuclei (STEM101; Takara, Y40400, IgG1, 1:100) and anti-human NOGOA (Santa Cruz Biotechnology, sc-11030, goat, 1:50) antibodies. In vivo characterization was performed using a series of primary antibodies listed in TableS2. For MBP staining, sections were pre-treated with ethanol (10min, RT). For glial-glial interactions, oligodendrocyte-specific connexin was detected with anti-Connexin 47 (Cx47; Invitrogen, 4A11A2, IgG1, 1:200) and astrocyte-specific connexin, with anti-Connexin 43 (Cx43; Sigma-Aldrich, C6219, rabbit, 1:50) and sections were pre-treated with methanol (10 min, -20°C). Secondary antibodies conjugated with FITC, TRITC (SouthernBiotech) or Alexa Fluor 647 (Life Technologies) were used respectively at 1:100 and 1:1000. Biotin-conjugated antibodies followed by AMCA AVIDIN D (Vector, A2006, 1:20). Nuclei were stained with Dapi (1 µg/ml, Sigma-Aldrich) (1:1000). Tissue scanning, cell visualization and imaging were performed with a Carl Zeiss microscope equipped with ApoTome 2.

#### Electron Microscopy

For electron microscopy, Shi/Shi:Rag2-/- mice grafted with control and RRMS hiOLs (n=4 per group) were perfused with 1% PBS followed by a mixture of 4% paraformaldehyde/5% glutaraldehyde (Electron Microscopy Science) in 1% PBS. After 2 hr post-fixation in the same solution, 100 µm-thick sagittal sections were cut and fixed in 2% osmium tetroxide (Sigma-Aldrich) overnight. After dehydration, samples were flat-embedded in Epon. Ultra-thin sections (80 nm) of the median corpus callosum were examined and imaged with a HITACHI 120kV HT-7700 electron microscope.

### Electrophysiology in vivo

Electrophysiological recordings were performed in mice grafted with MS- and control-hiOLs, and compared with non-grafted intact or medium injected Shi/Shi:Rag2–/– mice and wild-type mice 16 weeks after injection (n = 4–6 per group) as described (*13*). Briefly mice were anesthetized with 2 to 4% isoflurane performed under analgesia (0,1mg/kg buprecare) and placed in a stereotaxic frame (D. Kopf, Tujunga, CA, USA). Body temperature was maintained at 37°C by a feedback-controlled heating blanket (CMA Microdialysis). Electrical stimulation (0.1 ms at 0 – 0.1mA, was applied using a bipolar electrode (FHC-CBBSE75) inserted to a depth of 200 μm into the left cortex at 2 mm posterior to bregma and 3 mm from the midline. At the same coordinates in the contralateral hemisphere, home-made electrodes were positioned for recording local field potentials (LFPs) generated by transcallosal electric stimulation. Electrical stimulation and evoked LFPs were performed by the data acquisition system apparatus (Neurosoft, Russia) and signals were filtered at 0.01-1 000 Hz. Each response latency (in ms) was measured as the time between the onset of stimulus artifact to the first peak for each animal. A ground electrode was placed subcutaneously over the neck.

### Electrophysiology in acute brain slices

Slice preparation and recordings. Acute coronal slices (300µm) containing corpus callosum were made from Shi/Shi:Rag2-/- mice grafted with control (n = 7) and RRMS (n=6) RFP+ hiOLs. They were prepared from grafted mice between 12 and 15 wpg as previously described (*71*). Briefly, slices (100µm) were performed in a chilled cutting solution containing (in mM): 93 NMDG, 2.5 KCl, 1.2 NaH2PO4, 30 NaHCO3, 20 HEPES, 25 Glucose, 2 urea, 5 Na-ascorbate, 3 Na-pyruvate, 0.5 CaCl2, 10 MgCl2 (pH to 7.3-7.4; 95% O2, 5% CO2) and kept in the same solution for 8 min at 34°C. Then, they were transferred for 20 min to solution at 34°C containing (in mM): 126 NaCl, 2.5 KCl, 1.25 NaH2PO4, 26 NaHCO3, 20 Glucose, 5 Na-pyruvate, 2 CaCl2, 1 MgCl2 (pH to 7.3-7.4; 95% O2, 5% CO2). Transplanted RFP+ hiOLs were visualized with a 40x fluorescent water-immersion objective on an Olympus BX51 microscope coupled to a CMOS digital camera (TH4-200 OptiMOS) and an LED light source (CoolLed p-E2, Scientifica, UK), and recorded in voltage-clamp mode with an intracellular solution containing (in mM): 130 k-gluconate (KGlu), 0.1 EGTA, 2 MgCl2, 10 HEPES, 10 GABA, 2 Na2-ATP, 0.5 Na-GTP, 10 Na2-phosphocreatine, and biocytin 5.4 (pH ≈ 7.23). Holding potentials were corrected by a junction potential of -10 mV. Electrophysiological recordings were performed with Multiclamp 700B and Pclamp10.6 software (Molecular Devices). Signals were filtered at 3 kHz, digitized at 10 kHz and analyzed off-line.

Immunostainings and imaging of recorded slices. For analysis of recorded cells, one single RFP+ cell per hemisphere was recorded in a slice and loaded with biocytin for 25 min in whole-cell configuration. After gently removing the patch pipette, biocytin was allowed to diffuse for at least 10 min before the slice was fixed 2 h in 4% paraformaldehyde at 4°C. Then, the slice was rinsed three times in PBS for 10 min and incubated with 1% Triton X-100 and 10% normal goat serum (NGS) for 2 hr. After washing in PBS, slices were immunostained for SOX10, CC1, and STEM101/121. Tissues were incubated with primary antibodies for 3 days at 4°C. Secondary antibodies were diluted in 2% NGS and 0.2% Triton X-100. Tissues were incubated with secondary antibodies for 2 hr at room temperature. Biocytin was revealed with secondary antibodies using DyLight-488 streptavidin (Vector Labs, Burlingame, USA, 1:200). Images of biocytin-loaded cells were acquired either with a Carl Zeiss microscope equipped with ApoTome 2, or a LEICA SP8 confocal microscope (63X oil objective; NA=1.4; 0.75 µm Z-step) and processed with NIH ImageJ software (*72*).

### Automated quantification of oligodendrocyte myelination in vivo

We adapted the heuristic algorithm from (*32*) to identify STEM+MBP+ OLs in tissue sections. The foundations of the quantitative method remained the same. A ridge-filter extracted sheath-like objects based on intensity and segments associated to cell bodies using watershed segmentation. Two additional features adapted the workflow beyond its original in vitro application. First, we added functionality to allow colocalization of multiple fluorescent stains, as we needed to quantify triple positive STEM+/MBP+/DAPI+ cell objects. Second, since oligodendrocyte sheaths are not parallel and aligned in situ as they are in dissociated nanofiber cell cultures, we adapted the algorithm to report additional metrics about MBP production locally and globally that do not rely on the dissociation of sheaths in dense regions.

Cell nuclei were identified using watershed segmentation of DAPI+ regions and then colocalized pixel-wise with STEM+ objects. The DAPI+ nuclei were then used as local minima to seed a watershed segmentation of the STEM+ stain to separate nearby cell bodies. Finally, the identified STEM+ cell bodies were colocalized with overlapping MBP+ sheath-like ridges to define ensheathed cells. We reported the area of MBP overlapping with STEM fluorescence in colocalized regions associated with individual cells, as well as the number of single, double, and triple-fluorescently labelled cells. Additionally, different cellular phenotypes were noted in situ that were then captured with the adapted algorithm. Qualitatively, we observed cells with expansive MBP production without extended linear sheath-like segments that were not observed in previous applications of the algorithm. These cells were denoted as “tuft” cells, and were quantitatively defined as STEM+/MBP+/DAPI+ cells without fluorescent ridges that could be identified as extended sheath-like objects.

The myelination potential of 3 control and 3 MS lines was evaluated at 4, 8, 12, 16 and 20 wpg (n = 2-6 per line and per time point, n= 6-14 per time point). For each animal, 6 serial sections at 180 µm intervals were analyzed. The percentage of MBP+ cells (composed of ensheathed or tuft cells) was evaluated. Total MBP+ area per STEM+ cells and the average length of MBP+ sheaths per MBP+ cells were analyzed.

### Other quantification

#### Cell survival, proliferation and differentiation in vivo

The number of STEM101+ grafted cells expressing Caspase3, or Ki67, or SOX10 and CC1 was quantified in the core of the corpus callosum at 8, 12 and 16 wpg. For each animal (n=3 per group) 6 serial sections at 180 µm intervals were analyzed. Cell counts were expressed as the percentage of total STEM101+ cells.

#### Myelination by electron microscopy

G-ratio (diameter of axon/diameter of axon and myelin sheath) of donor-derived compact myelin was measured as previously described (*12*). Briefly, the maximum and minimum diameters of a given axon and the maximum and minimum axon plus myelin sheath diameter were measured with the ImageJ software at a magnification of 62 000 for a minimum of 70 myelinated axons per animal. Data were expressed as the mean of the maximal and minimal values for each axon for mice from each group (n=4 mice/group). Myelin compaction was confirmed at a magnification of 220 000.

### Statistical analysis

Data are presented as mean + SEM. Statistical significance was determined by two-tailed Mann Whitney U test when comparing two statistical groups, and with one-way or two-way ANOVA followed by Tukey’s or Dunnett’s (in vivo electrophysiology) multiple comparison tests for multiple groups. Since electrophysiological data in brain slices do not follow a normal distribution after a D’Agostino-Pearson normality test, we also performed two-tailed Mann-Whitney U test for comparison between groups. Statistics were done in GraphPad Prism 5.00 and GraphPad Prism 8.2.1 (GraphPad Software Inc, USA). See the figure captions for the test used in each experiment.

## Author contributions

Conceptualization, S.M. and A.B.V.E.; Methodology, S.M., L.S., B.M., T.X., B.G.D, M.L.S., D.R., L.O., K-P.K, H.S., J.A., T.K., G.M., T.K., M.C.A. and A.B.V.E.; Formal Analysis, S.M., B.M-S., T.X., M.C.A., A.B.V.E.; Writing, S.M., A.B.V.E. with editing and discussion from all co-authors; Funding Acquisition, S.M., A.B.V.E.; Supervision, A.B.V.E.

## Data and materials availability

All data associated with this study are available in the main text or the supplementary materials

## Supplementary Materials

**Fig. S1.**
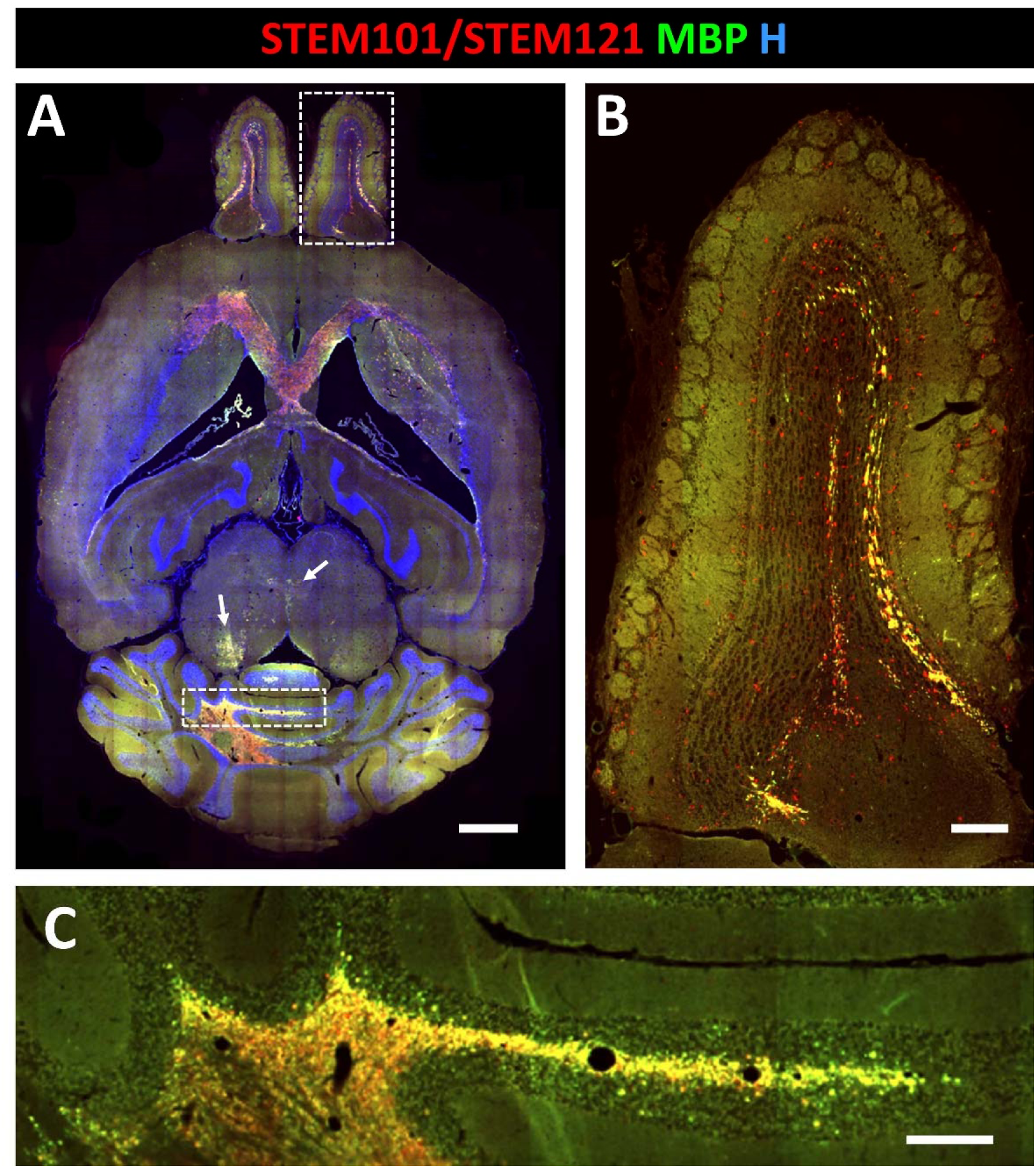
MS-hiOL derived progeny extensively myelinates the dysmyelinated Shi/Shi:Rag2-/- forebrain. (**A**) General view illustrating combined detection of human nuclei STEM101 and human cytoplasm STEM121 (red) with MBP (green) in horizontal sections of the *Shi/Shi Rag2*^*-/-*^ brain 20 wpg. Human cells and their myelin can be found rostrally in the olfactory bulb, medially in the corpus callosum (graft location) as well as caudally in the pons (arrows) and the cerebellum. (**B**) Enlargement of the vertical boxed area of the olfactory bulb. (**C**) Enlargement of the horizontal boxed area in the cerebellum. H: Hoechst dye; Scale bar: 1 mm in **a**, 200 μm in **B** and **C**.

**Fig. S2.**
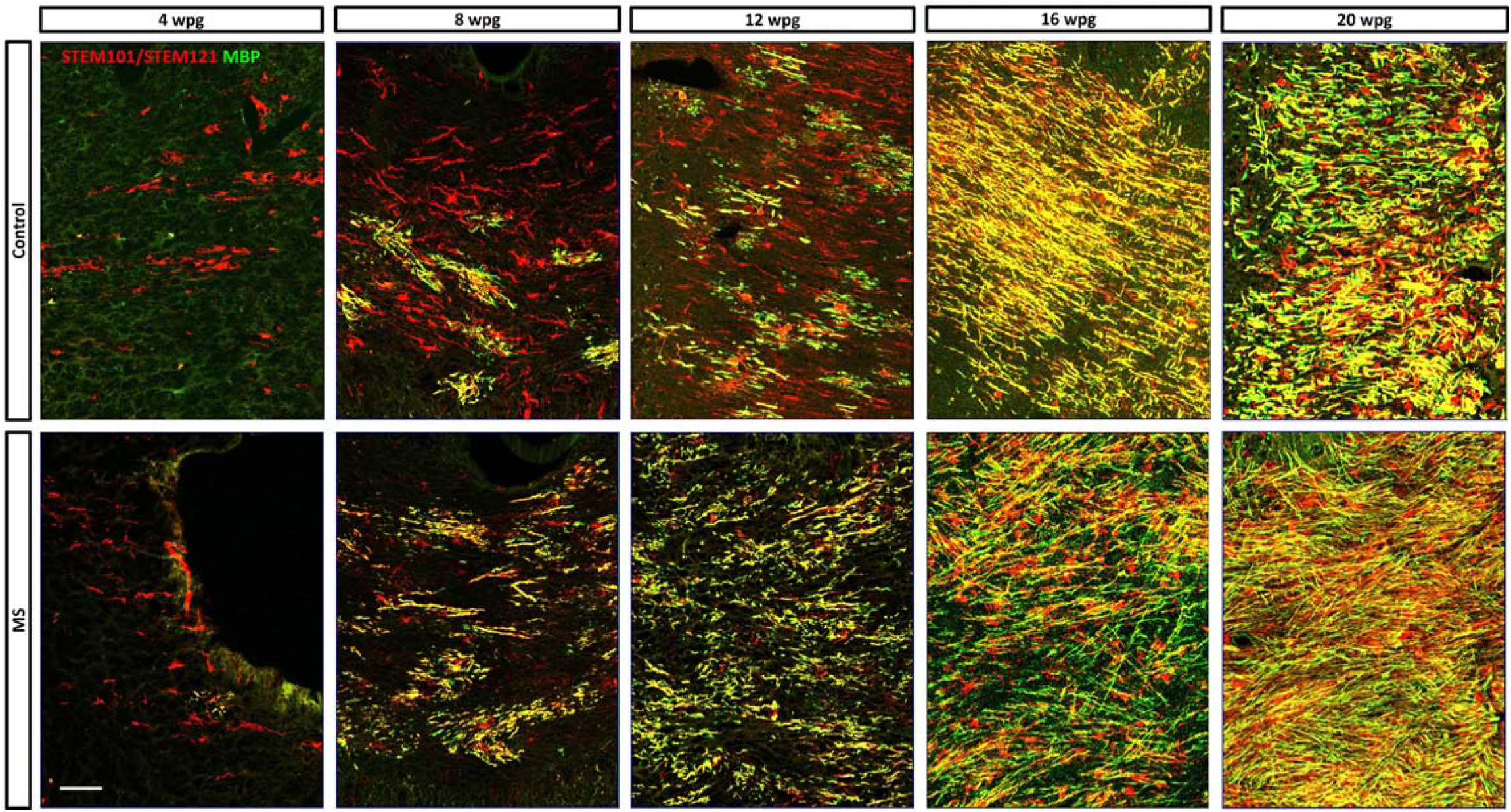
Myelination by the most potent MS and control hiOL lines. (**A**) Combined immunodetection of human nuclei STEM101 and human cytoplasm STEM121 (red) with MBP (green) in the *Shi/Shi Rag2*^*–/–*^ corpus callosum at 4, 8, 12, 16 and 20 wpg. Views of horizontal sections at the level of the corpus callosum showing the progressive increase of donor derived myelin for control- (top) and MS- (bottom) hiOLs. Wpg: weeks post graft. H, Hoechst dye. Scale bar: 50.

**Fig. S3.**
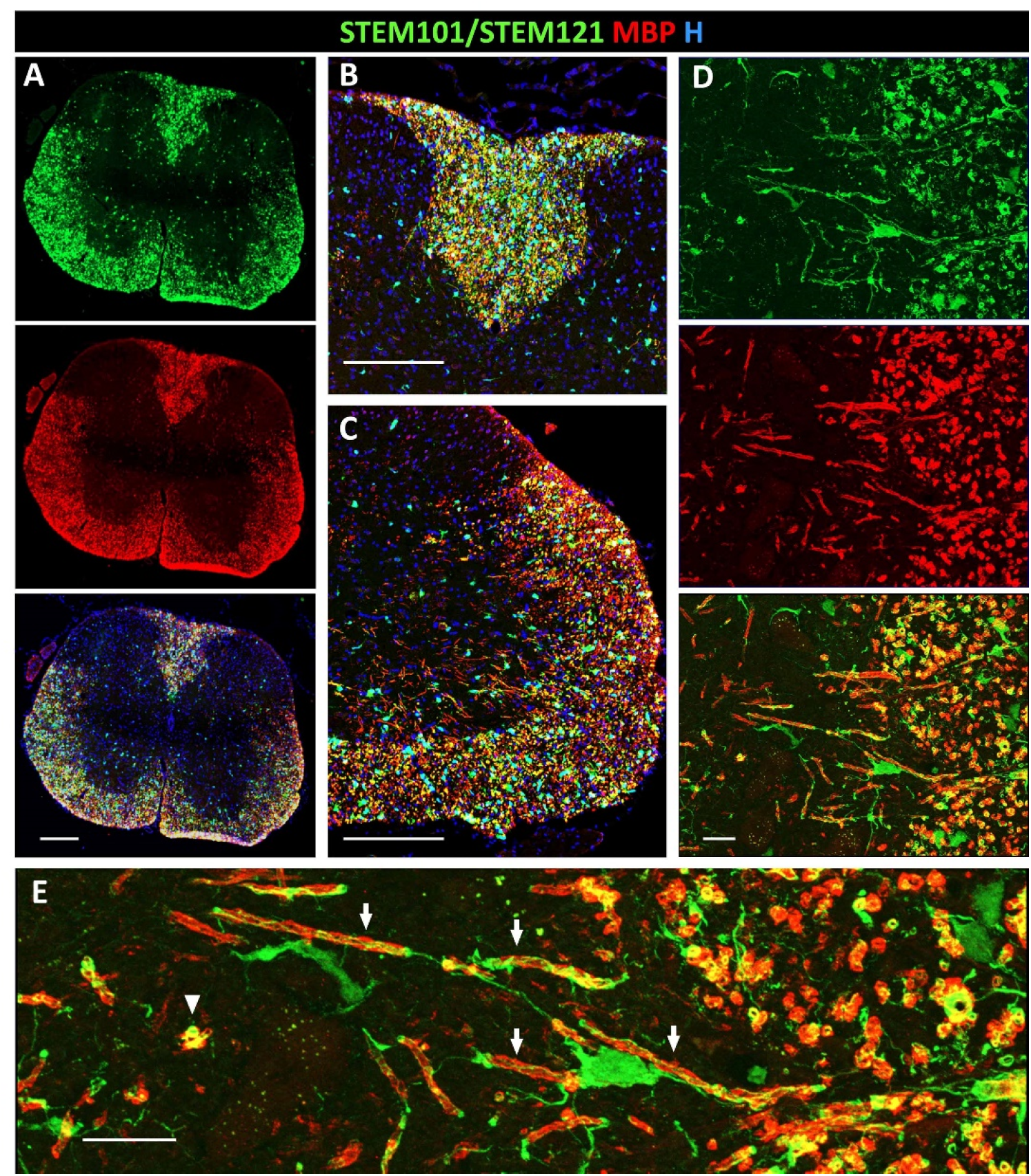
MS-hiOL-derived progeny extensively myelinate the dysmyelinated Shi/Shi: Rag2– /– spinal cord. MS-hiOL were grafted at the thoracic level of the post-natal spinal cord (4 weeks) and animals sacrificed 12 wpg. (**A**) General view of a spinal cord cross section illustrating the single (top and middle panels) and combined (lower panel) detection of human nuclei and human cytoplasmic antigen STEM101/121 (green) and MBP (red); STEM101/121 and MBP staining fully overlap in the dorsal funiculus where cells were grafted, but also in the ventral and lateral white matter where cells presumably migrated (**B** and **C**). Higher magnifications of the dorsal funiculus (**B**), and the ventro-lateral white matter (**C**) illustrating the overlapping staining. (**D**) Enlargement of (**C**) illustrating a human oligodendrocyte connected to several human-derived myelin internodes (arrows). Several donut-shaped STEM101/STEM121+/MBP+ structures corresponding to cross-sections of axons surrounded by human myelin are also observed (arrowhead points to such example); top and middle, single channels; bottom, combined channels. (**E**) Higher magnification of **D** illustrating the connections of the human oligodendrocyte cell processes to the myelin internodes. Note the presence of the STEM cytoplasmic marker in the non-compacted parts of myelin (paranodes and outer loops). H: Hoechst dye; Scale bar: 200 μm in A-C, 20 μm in **D** and **E**.

**Fig. S4.**
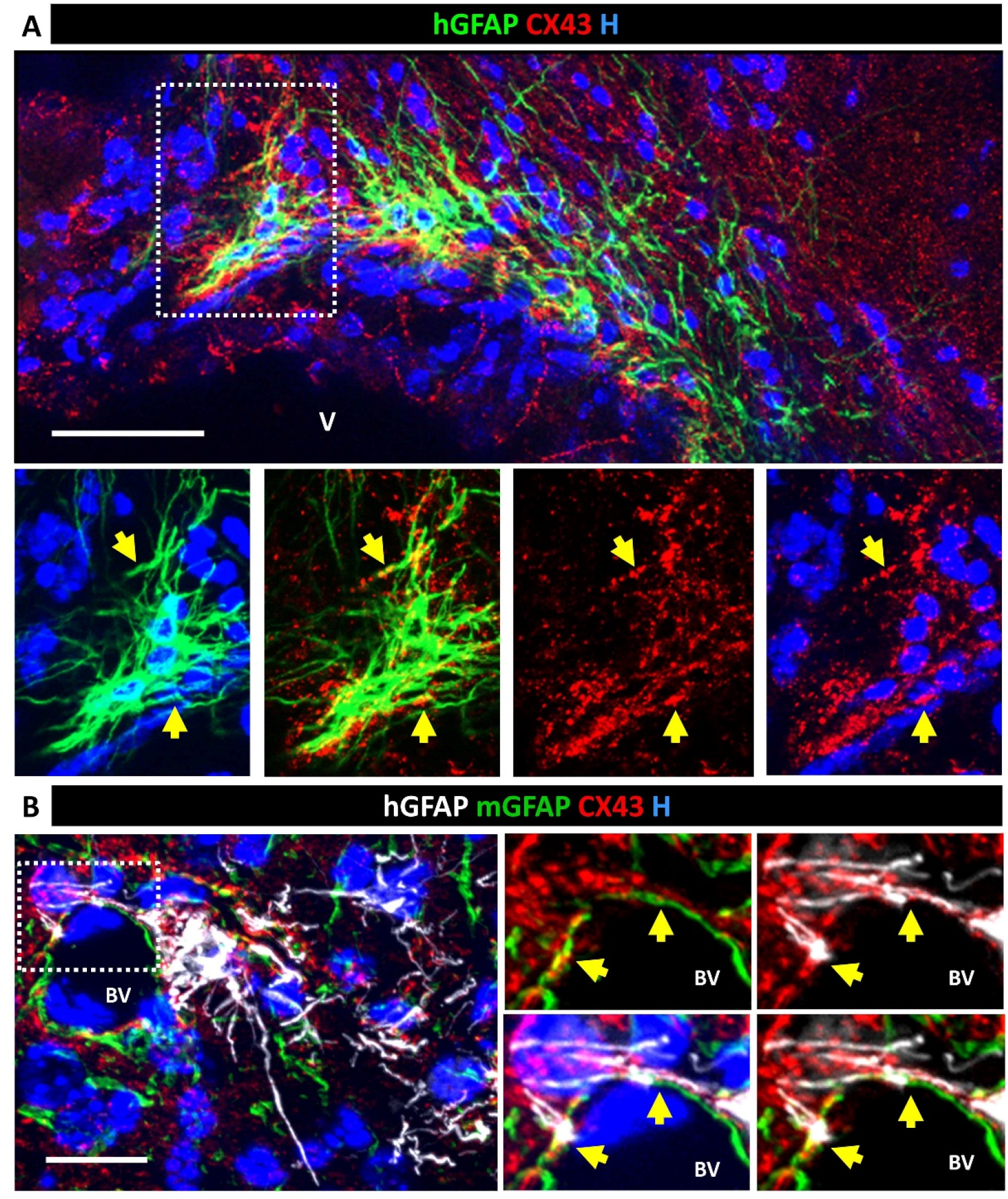
Grafted hiOL-derived astrocytes express astrocyte-gap junction plaque Cx43. (**A**) Immunodetection of hGFAP (green), and astrocyte Cx43 illustrates in the top panel, the presence of a group of MS-hiOL-derived astrocytes nested in the subventricular zone of the lateral ventricle (V); the boxed area enlarged in the lower panels, shows Cx47+ gap junction plaques decorating the human astrocyte cell body and processes (yellow arrows). (**B**) Combined immunodetection of human GFAP (white), mouse GFAP (green) and astrocyte Cx43, illustrates the integration of human astrocytes around blood vessels (BV) and the presence of Cx43+ gap junction plaques between human and murine perivascular astrocyte processes. The boxed area in the left panel is enlarged in the right panel and highlights Cx43+ gap junction plaques shared by the murine and human astrocyte processes (yellow arrows). H: Hoechst dye; scale bars: 50 µm for **A**, 20 µm for **A**.

**Table S1.** Overview of cell lines

| Cell line | Group | Age at donation | Gender | Disease duration at timepoint of biopsy | Relationship |
| --- | --- | --- | --- | --- | --- |
| C1 | Control | 32 | F | na | Monozygous twin |
| C2 | Control | newborn | M | na | Non relative |
| C3 | Control | 34 | F | na | Monozygous twin |
| RRMS1 | RRMS | 32 | F | 11 | Monozygous twin |
| RRMS2 | RRMS | 36 | M | 12 | Non relative |
| RRMS3 | RRMS | 34 | F | 3 | Monozygous twin |

**Table S2.** Primary antibodies

| Antibody | Dilution | Source |
| --- | --- | --- |
| Rabbit Anti-Ki67 | 1:100 | abcam, ab15580 |
| Goat anti-hNOGOA | 1:50 | Santa Cruz Biotechnology, sc-11030 |
| Mouse anti human nuclei, STEM101 | 1:100 | Takara, Y40400 |
| Mouse anti human cytoplasm, STEM121 | 1:100 | Takara, Y40410 |
| Rat anti-MBP | 1:100 | abcam, ab7349 |
| Mouse anti-MBP | 1:100 | Millipore, MAB387 |
| Mouse anti-MBP | 1:500 | Novus Biologicals, NBP2-22121 |
| Rabbit anti-OLIG2 | 1:400 | Millipore, AB 9610 |
| Mouse anti-OLIG2 | 1:500 | Sigma-Aldrich, MABN50 |
| Mouse anti-CC1 | 1:100 | Millipore, OP80 |
| Goat anti-SOX10 | 1:50 | R&D systems, AF2864 |
| Rabbit anti-NF-200 | 1:200 | Sigma-Aldrich, N4142 |
| Mouse anti human GFAP, STEM123 | 1:500 | Takara, Y40420 |
| Goat anti GFAP | 1:200 | Thermo-Fisher Scientific, PA5-18598 |
| Rabbit anti-Caspase 3 | 1:100 | Cell signaling, 9661 |
| Rabbit anti-Caspr | 1:500 | abcam, ab34151 |
| Mouse anti-AnkyrinG | 1:100 | Santa Cruz Biotech<br>Sc-166602 |
| Rabbit anti-Neurofascin | 1:100 | abcam, ab31457 |
| Mouse anti-Cx47 | 1:200 | Invitrogen, 4A11A2 |
| Rabbit anti-Cx43 | 1:50 | Sigma-Aldrich, C6219 |

## References

1. T. Goldschmidt, J. Antel, F. B. Konig, W. Bruck, T. Kuhlmann, Remyelination capacity of the MS brain decreases with disease chronicity. Neurology 72, 1914–1921 (2009).

2. J. T. Chen, D. L. Collins, H. L. Atkins, M. S. Freedman, D. L. Arnold, M. S. B. M. T. S. G. Canadian, Magnetization transfer ratio evolution with demyelination and remyelination in multiple sclerosis lesions. Ann Neurol 63, 254–262 (2008).

3. P. Patrikios, C. Stadelmann, A. Kutzelnigg, H. Rauschka, M. Schmidbauer, H. Laursen, P. S. Sorensen, W. Bruck, C. Lucchinetti, H. Lassmann, Remyelination is extensive in a subset of multiple sclerosis patients. Brain 129, 3165–3172 (2006).

4. R. Patani, M. Balaratnam, A. Vora, R. Reynolds, Remyelination can be extensive in multiple sclerosis despite a long disease course. Neuropathol Appl Neurobiol 33, 277–287 (2007).

5. B. Bodini, M. Veronese, D. Garcia-Lorenzo, M. Battaglini, E. Poirion, A. Chardain, L. Freeman, C. Louapre, M. Tchikviladze, C. Papeix, F. Dolle, B. Zalc, C. Lubetzki, M. Bottlaender, F. Turkheimer, B. Stankoff, Dynamic Imaging of Individual Remyelination Profiles in Multiple Sclerosis. Ann Neurol 79, 726–738 (2016).

6. N. Ohno, K. Ikenaka, Axonal and neuronal degeneration in myelin diseases. Neurosci Res 139, 48–57 (2019).

7. A. Chang, W. W. Tourtellotte, R. Rudick, B. D. Trapp, Premyelinating oligodendrocytes in chronic lesions of multiple sclerosis. N Engl J Med 346, 165–173 (2002).

8. T. Kuhlmann, V. Miron, Q. Cui, C. Wegner, J. Antel, W. Bruck, Differentiation block of oligodendroglial progenitor cells as a cause for remyelination failure in chronic multiple sclerosis. Brain 131, 1749–1758 (2008).

9. G. Wolswijk, Chronic stage multiple sclerosis lesions contain a relatively quiescent population of oligodendrocyte precursor cells. J Neurosci 18, 601–609 (1998).

10. S. Mozafari, M. A. Sherafat, M. Javan, J. Mirnajafi-Zadeh, T. Tiraihi, Visual evoked potentials and MBP gene expression imply endogenous myelin repair in adult rat optic nerve and chiasm following local lysolecithin induced demyelination. Brain Res 1351, 50–56 (2010).

11. K. J. Smith, W. I. McDonald, W. F. Blakemore, Restoration of secure conduction by central demyelination. Trans Am Neurol Assoc 104, 25–29 (1979).

12. S. Mozafari, C. Laterza, D. Roussel, C. Bachelin, A. Marteyn, C. Deboux, G. Martino, A. Baron-Van Evercooren, Skin-derived neural precursors competitively generate functional myelin in adult demyelinated mice. J Clin Invest 125, 3642–3656 (2015).

13. M. S. Windrem, S. J. Schanz, M. Guo, G. F. Tian, V. Washco, N. Stanwood, M. Rasband, N. S. Roy, M. Nedergaard, L. A. Havton, S. Wang, S. A. Goldman, Neonatal chimerization with human glial progenitor cells can both remyelinate and rescue the otherwise lethally hypomyelinated shiverer mouse. Cell Stem Cell 2, 553–565 (2008).

14. C. Laterza, A. Merlini, D. De Feo, F. Ruffini, R. Menon, M. Onorati, E. Fredrickx, L. Muzio, A. Lombardo, G. Comi, A. Quattrini, C. Taveggia, C. Farina, E. Cattaneo, G. Martino, iPSC-derived neural precursors exert a neuroprotective role in immune-mediated demyelination via the secretion of LIF. Nat Commun 4, 2597 (2013).

15. F. Mei, K. Lehmann-Horn, Y. A. Shen, K. A. Rankin, K. J. Stebbins, D. S. Lorrain, K. Pekarek, A. S. S, L. Xiao, C. Teuscher, H. C. von Budingen, J. Wess, J. J. Lawrence, A. J. Green, S. P. Fancy, S. S. Zamvil, J. R. Chan, Accelerated remyelination during inflammatory demyelination prevents axonal loss and improves functional recovery. Elife 5, (2016).

16. D. Verden, W. B. Macklin, Neuroprotection by central nervous system remyelination: Molecular, cellular, and functional considerations. J Neurosci Res 94, 1411–1420 (2016).

17. S. Y. Leong, V. T. Rao, J. M. Bin, P. Gris, M. Sangaralingam, T. E. Kennedy, J. P. Antel, Heterogeneity of oligodendrocyte progenitor cells in adult human brain. Ann Clin Transl Neurol 1, 272–283 (2014).

18. A. M. Falcao, D. van Bruggen, S. Marques, M. Meijer, S. Jakel, E. Agirre, Samudyata, E. M. Floriddia, D. P. Vanichkina, C. Ffrench-Constant, A. Williams, A. O. Guerreiro-Cacais, G. Castelo-Branco, Disease-specific oligodendrocyte lineage cells arise in multiple sclerosis. Nat Med 24, 1837–1844 (2018).

19. S. Jakel, E. Agirre, A. Mendanha Falcao, D. van Bruggen, K. W. Lee, I. Knuesel, D. Malhotra, C. Ffrench-Constant, A. Williams, G. Castelo-Branco, Altered human oligodendrocyte heterogeneity in multiple sclerosis. Nature 566, 543–547 (2019).

20. M. Osipovitch, A. Asenjo Martinez, J. N. Mariani, A. Cornwell, S. Dhaliwal, L. Zou, D. Chandler-Militello, S. Wang, X. Li, S. J. Benraiss, R. Agate, A. Lampp, A. Benraiss, M. S. Windrem, S. A. Goldman, Human ESC-Derived Chimeric Mouse Models of Huntington’s Disease Reveal Cell-Intrinsic Defects in Glial Progenitor Cell Differentiation. Cell Stem Cell 24, 107–122 e107 (2019).

21. D. L. McPhie, R. Nehme, C. Ravichandran, S. M. Babb, S. D. Ghosh, A. Staskus, A. Kalinowski, R. Kaur, P. Douvaras, F. Du, D. Ongur, V. Fossati, K. Eggan, B. M. Cohen, Oligodendrocyte differentiation of induced pluripotent stem cells derived from subjects with schizophrenias implicate abnormalities in development. Transl Psychiatry 8, 230 (2018).

22. M. S. Windrem, M. Osipovitch, Z. Liu, J. Bates, D. Chandler-Militello, L. Zou, J. Munir, S. Schanz, K. McCoy, R. H. Miller, S. Wang, M. Nedergaard, R. L. Findling, P. J. Tesar, S. A. Goldman, Human iPSC Glial Mouse Chimeras Reveal Glial Contributions to Schizophrenia. Cell Stem Cell 21, 195–208 e196 (2017).

23. A. M. Nicaise, E. Banda, R. M. Guzzo, K. Russomanno, W. Castro-Borrero, C. M. Willis, K. M. Johnson, A. C. Lo, S. J. Crocker, iPS-derived neural progenitor cells from PPMS patients reveal defect in myelin injury response. Exp Neurol 288, 114–121 (2017).

24. A. M. Nicaise, L. J. Wagstaff, C. M. Willis, C. Paisie, H. Chandok, P. Robson, V. Fossati, A. Williams, S. J. Crocker, Cellular senescence in progenitor cells contributes to diminished remyelination potential in progressive multiple sclerosis. Proc Natl Acad Sci U S A 116, 9030–9039 (2019).

25. P. Douvaras, J. Wang, M. Zimmer, S. Hanchuk, M. A. O’Bara, S. Sadiq, F. J. Sim, J. Goldman, V. Fossati, Efficient generation of myelinating oligodendrocytes from primary progressive multiple sclerosis patients by induced pluripotent stem cells. Stem Cell Reports 3, 250–259 (2014).

26. J. A. Garcia-Leon, M. Kumar, R. Boon, D. Chau, J. One, E. Wolfs, K. Eggermont, P. Berckmans, N. Gunhanlar, F. de Vrij, B. Lendemeijer, B. Pavie, N. Corthout, S. A. Kushner, J. C. Davila, I. Lambrichts, W. S. Hu, C. M. Verfaillie, SOX10 Single Transcription Factor-Based Fast and Efficient Generation of Oligodendrocytes from Human Pluripotent Stem Cells. Stem Cell Reports 10, 655–672 (2018).

27. B. Song, G. Sun, D. Herszfeld, A. Sylvain, N. V. Campanale, C. E. Hirst, S. Caine, H. C. Parkington, M. A. Tonta, H. A. Coleman, M. Short, S. D. Ricardo, B. Reubinoff, C. C. Bernard, Neural differentiation of patient specific iPS cells as a novel approach to study the pathophysiology of multiple sclerosis. Stem Cell Res 8, 259–273 (2012).

28. M. Ehrlich, S. Mozafari, M. Glatza, L. Starost, S. Velychko, A. L. Hallmann, Q. L. Cui, A. Schambach, K. P. Kim, C. Bachelin, A. Marteyn, G. Hargus, R. M. Johnson, J. Antel, J. Sterneckert, H. Zaehres, H. R. Scholer, A. Baron-Van Evercooren, T. Kuhlmann, Rapid and efficient generation of oligodendrocytes from human induced pluripotent stem cells using transcription factors. Proc Natl Acad Sci U S A 114, E2243–E2252 (2017).

29. D. Seilhean, A. Gansmuller, A. Baron-Van Evercooren, M. Gumpel, F. Lachapelle, Myelination by transplanted human and mouse central nervous system tissue after long-term cryopreservation. Acta Neuropathol 91, 82–88 (1996).

30. M. S. Windrem, M. C. Nunes, W. K. Rashbaum, T. H. Schwartz, R. A. Goodman, G. McKhann, 2nd, N. S. Roy, S. A. Goldman, Fetal and adult human oligodendrocyte progenitor cell isolates myelinate the congenitally dysmyelinated brain. Nat Med 10, 93–97 (2004).

31. S. Wang, J. Bates, X. Li, S. Schanz, D. Chandler-Militello, C. Levine, N. Maherali, L. Studer, K. Hochedlinger, M. Windrem, S. A. Goldman, Human iPSC-derived oligodendrocyte progenitor cells can myelinate and rescue a mouse model of congenital hypomyelination. Cell Stem Cell 12, 252–264 (2013).

32. Y. K. T. Xu, D. Chitsaz, R. A. Brown, Q. L. Cui, M. A. Dabarno, J. P. Antel, T. E. Kennedy, Deep learning for high-throughput quantification of oligodendrocyte ensheathment at single-cell resolution. Commun Biol 2, 116 (2019).

33. M. E. Bechler, L. Byrne, C. Ffrench-Constant, CNS Myelin Sheath Lengths Are an Intrinsic Property of Oligodendrocytes. Curr Biol 25, 2411–2416 (2015).

34. A. Gansmuller, F. Lachapelle, A. Baron-Van Evercooren, J. J. Hauw, N. Baumann, M. Gumpel, Transplantations of newborn CNS fragments into the brain of shiverer mutant mice: extensive myelination by transplanted oligodendrocytes. II. Electron microscopic study. Dev Neurosci 8, 197–207 (1986).

35. C. A. Ruff, H. Ye, J. M. Legasto, N. A. Stribbell, J. Wang, L. Zhang, M. G. Fehlings, Effects of adult neural precursor-derived myelination on axonal function in the perinatal congenitally dysmyelinated brain: optimizing time of intervention, developing accurate prediction models, and enhancing performance. J Neurosci 33, 11899–11915 (2013).

36. M. Kukley, A. Nishiyama, D. Dietrich, The fate of synaptic input to NG2 glial cells: neurons specifically downregulate transmitter release onto differentiating oligodendroglial cells. J Neurosci 30, 8320–8331 (2010).

37. A. Sahel, F. C. Ortiz, C. Kerninon, P. P. Maldonado, M. C. Angulo, B. Nait-Oumesmar, Alteration of synaptic connectivity of oligodendrocyte precursor cells following demyelination. Front Cell Neurosci 9, 77 (2015).

38. M. Maglione, O. Tress, B. Haas, K. Karram, J. Trotter, K. Willecke, H. Kettenmann, Oligodendrocytes in mouse corpus callosum are coupled via gap junction channels formed by connexin47 and connexin32. Glia 58, 1104–1117 (2010).

39. A. Meyer, S. C. Yadav, K. Dedek, Phenotyping of Gap-Junctional Coupling in the Mouse Retina. Methods Mol Biol 1753, 249–259 (2018).

40. R. Basu, J. D. Sarma, Connexin 43/47 channels are important for astrocyte/ oligodendrocyte cross-talk in myelination and demyelination. J Biosci 43, 1055–1068 (2018).

41. C. Papaneophytou, E. Georgiou, K. A. Kleopa, The role of oligodendrocyte gap junctions in neuroinflammation. Channels (Austin) 13, 247–263 (2019).

42. C. Vilarino-Guell, A. Zimprich, F. Martinelli-Boneschi, B. Herculano, Z. Wang, F. Matesanz, E. Urcelay, K. Vandenbroeck, L. Leyva, D. Gris, C. Massaad, J. A. Quandt, A. L. Traboulsee, M. Encarnacion, C. Q. Bernales, J. Follett, I. M. Yee, M. G. Criscuoli, A. Deutschlander, E. M. Reinthaler, T. Zrzavy, E. Mascia, A. Zauli, F. Esposito, A. Alcina, G. Izquierdo, L. Espino-Paisan, J. Mena, A. Antiguedad, P. Urbaneja-Romero, J. Ortega-Pinazo, W. Song, A. D. Sadovnick, Exome sequencing in multiple sclerosis families identifies 12 candidate genes and nominates biological pathways for the genesis of disease. PLoS Genet 15, e1008180 (2019).

43. R. J. M. Franklin, C. Ffrench-Constant, Regenerating CNS myelin - from mechanisms to experimental medicines. Nat Rev Neurosci 18, 753–769 (2017).

44. M. Stangel, T. Kuhlmann, P. M. Matthews, T. J. Kilpatrick, Achievements and obstacles of remyelinating therapies in multiple sclerosis. Nat Rev Neurol 13, 742–754 (2017).

45. G. J. Duncan, S. B. Manesh, B. J. Hilton, P. Assinck, J. Liu, A. Moulson, J. R. Plemel, W. Tetzlaff, Locomotor recovery following contusive spinal cord injury does not require oligodendrocyte remyelination. Nat Commun 9, 3066 (2018).

46. M. S. Y. Yeung, M. Djelloul, E. Steiner, S. Bernard, M. Salehpour, G. Possnert, L. Brundin, J. Frisen, Dynamics of oligodendrocyte generation in multiple sclerosis. Nature 566, 538–542 (2019).

47. K. Chanoumidou, S. Mozafari, A. Baron-Van Evercooren, T. Kuhlmann, Stem cell derived oligodendrocytes to study myelin diseases. Glia 68, 705–720 (2020).

48. F. J. Najm, A. M. Lager, A. Zaremba, K. Wyatt, A. V. Caprariello, D. C. Factor, R. T. Karl, T. Maeda, R. H. Miller, P. J. Tesar, Transcription factor-mediated reprogramming of fibroblasts to expandable, myelinogenic oligodendrocyte progenitor cells. Nat. Biotechnol 31, 426–433 (2013).

49. N. Yang, J. B. Zuchero, H. Ahlenius, S. Marro, Y. H. Ng, T. Vierbuchen, J. S. Hawkins, R. Geissler, B. A. Barres, M. Wernig, Generation of oligodendroglial cells by direct lineage conversion. Nat. Biotechnol 31, 434–439 (2013).

50. M. R. Livesey, D. Magnani, E. M. Cleary, N. A. Vasistha, O. T. James, B. T. Selvaraj, K. Burr, D. Story, C. E. Shaw, P. C. Kind, G. E. Hardingham, D. J. Wyllie, S. Chandran, Maturation and electrophysiological properties of human pluripotent stem cell-derived oligodendrocytes. Stem Cells 34, 1040–1053 (2016).

51. O. Tress, M. Maglione, D. May, T. Pivneva, N. Richter, J. Seyfarth, S. Binder, A. Zlomuzica, G. Seifert, M. Theis, E. Dere, H. Kettenmann, K. Willecke, Panglial gap junctional communication is essential for maintenance of myelin in the CNS. J Neurosci 32, 7499–7518 (2012).

52. T. Li, C. Giaume, L. Xiao, Connexins-mediated glia networking impacts myelination and remyelination in the central nervous system. Mol Neurobiol 49, 1460–1471 (2014).

53. N. Kamasawa, A. Sik, M. Morita, T. Yasumura, K. G. Davidson, J. I. Nagy, J. E. Rash, Connexin-47 and connexin-32 in gap junctions of oligodendrocyte somata, myelin sheaths, paranodal loops and Schmidt-Lanterman incisures: implications for ionic homeostasis and potassium siphoning. Neuroscience 136, 65–86 (2005).

54. S. Y. Chong, S. S. Rosenberg, S. P. Fancy, C. Zhao, Y. A. Shen, A. T. Hahn, A. W. McGee, X. Xu, B. Zheng, L. I. Zhang, D. H. Rowitch, R. J. Franklin, Q. R. Lu, J. R. Chan, Neurite outgrowth inhibitor Nogo-A establishes spatial segregation and extent of oligodendrocyte myelination. Proc Natl Acad Sci U S A 109, 1299–1304 (2012).

55. S. S. Rosenberg, E. E. Kelland, E. Tokar, A. R. De la Torre, J. R. Chan, The geometric and spatial constraints of the microenvironment induce oligodendrocyte differentiation. Proc Natl Acad Sci U S A 105, 14662–14667 (2008).

56. S. Vejar, J. E. Oyarzun, M. A. Retamal, F. C. Ortiz, J. A. Orellana, Connexin and Pannexin-Based Channels in Oligodendrocytes: Implications in Brain Health and Disease. Front Cell Neurosci 13, 3 (2019).

57. K. Masaki, Early disruption of glial communication via connexin gap junction in multiple sclerosis, Balo’s disease and neuromyelitis optica. Neuropathology 35, 469–480 (2015).

58. K. Masaki, S. O. Suzuki, T. Matsushita, T. Matsuoka, S. Imamura, R. Yamasaki, M. Suzuki, T. Suenaga, T. Iwaki, J. Kira, Connexin 43 astrocytopathy linked to rapidly progressive multiple sclerosis and neuromyelitis optica. PLoS One 8, e72919 (2013).

59. J. E. Rash, Molecular disruptions of the panglial syncytium block potassium siphoning and axonal saltatory conduction: pertinence to neuromyelitis optica and other demyelinating diseases of the central nervous system. Neuroscience 168, 982–1008 (2010).

60. K. A. Kleopa, S. S. Scherer, Molecular genetics of X-linked Charcot-Marie-Tooth disease. Neuromolecular Med 8, 107–122 (2006).

61. B. Uhlenberg, M. Schuelke, F. Ruschendorf, N. Ruf, A. M. Kaindl, M. Henneke, H. Thiele, G. Stoltenburg-Didinger, F. Aksu, H. Topaloglu, P. Nurnberg, C. Hubner, B. Weschke, J. Gartner, Mutations in the gene encoding gap junction protein alpha 12 (connexin 46.6) cause Pelizaeus-Merzbacher-like disease. Am J Hum Genet 75, 251–260 (2004).

62. B. Odermatt, K. Wellershaus, A. Wallraff, G. Seifert, J. Degen, C. Euwens, B. Fuss, H. Bussow, K. Schilling, C. Steinhauser, K. Willecke, Connexin 47 (Cx47)-deficient mice with enhanced green fluorescent protein reporter gene reveal predominant oligodendrocytic expression of Cx47 and display vacuolized myelin in the CNS. J Neurosci 23, 4549–4559 (2003).

63. X. Han, M. Chen, F. Wang, M. Windrem, S. Wang, S. Shanz, Q. Xu, N. A. Oberheim, L. Bekar, S. Betstadt, A. J. Silva, T. Takano, S. A. Goldman, M. Nedergaard, Forebrain engraftment by human glial progenitor cells enhances synaptic plasticity and learning in adult mice. Cell Stem Cell 12, 342–353 (2013).

64. C. International Multiple Sclerosis Genetics, Multiple sclerosis genomic map implicates peripheral immune cells and microglia in susceptibility. Science 365, (2019).

65. L. Kirby, J. Jin, J. G. Cardona, M. D. Smith, K. A. Martin, J. Wang, H. Strasburger, L. Herbst, M. Alexis, J. Karnell, T. Davidson, R. Dutta, J. Goverman, D. Bergles, P. A. Calabresi, Oligodendrocyte precursor cells present antigen and are cytotoxic targets in inflammatory demyelination. Nat Commun 10, 3887 (2019).

66. M. El Behi, C. Sanson, C. Bachelin, L. Guillot-Noel, J. Fransson, B. Stankoff, E. Maillart, N. Sarrazin, V. Guillemot, H. Abdi, I. Cournu-Rebeix, B. Fontaine, V. Zujovic, Adaptive human immunity drives remyelination in a mouse model of demyelination. Brain 140, 967–980 (2017).

67. P. Reinhardt, M. Glatza, K. Hemmer, Y. Tsytsyura, C. S. Thiel, S. Hoing, S. Moritz, J. A. Parga, L. Wagner, J. M. Bruder, G. Wu, B. Schmid, A. Ropke, J. Klingauf, J. C. Schwamborn, T. Gasser, H. R. Scholer, J. Sterneckert, Derivation and expansion using only small molecules of human neural progenitors for neurodegenerative disease modeling. PLoS. One 8, e59252 (2013).

68. M. Ehrlich, A. L. Hallmann, P. Reinhardt, M. J. rauzo-Bravo, S. Korr, A. Ropke, O. E. Psathaki, P. Ehling, S. G. Meuth, A. L. Oblak, J. R. Murrell, B. Ghetti, H. Zaehres, H. R. Scholer, J. Sterneckert, T. Kuhlmann, G. Hargus, Distinct Neurodegenerative Changes in an Induced Pluripotent Stem Cell Model of Frontotemporal Dementia Linked to Mutant TAU Protein. Stem Cell Reports 5, 83–96 (2015).

69. F. Lachapelle, E. Duhamel-Clerin, A. Gansmuller, A. Baron-Van Evercooren, H. Villarroya, M. Gumpel, Transplanted transgenically marked oligodendrocytes survive, migrate and myelinate in the normal mouse brain as they do in the shiverer mouse brain. Eur J Neurosci 6, 814–824 (1994).

70. A. Marteyn, N. Sarrazin, J. Yan, C. Bachelin, C. Deboux, M. D. Santin, P. Gressens, V. Zujovic, A. Baron-Van Evercooren, Modulation of the Innate Immune Response by Human Neural Precursors Prevails over Oligodendrocyte Progenitor Remyelination to Rescue a Severe Model of Pelizaeus-Merzbacher Disease. Stem Cells 34, 984–996 (2016).

71. F. C. Ortiz, C. Habermacher, M. Graciarena, P. Y. Houry, A. Nishiyama, B. Nait Oumesmar, M. C. Angulo, Neuronal activity in vivo enhances functional myelin repair. JCI Insight 5, (2019).

72. C. A. Schneider, W. S. Rasband, K. W. Eliceiri, NIH Image to ImageJ: 25 years of image analysis. Nat Methods 9, 671–675 (2012).

